# A common neural code for meaning in discourse production and comprehension

**DOI:** 10.1101/2022.10.15.512349

**Authors:** Tanvi Patel, Matías Morales, Martin J. Pickering, Paul Hoffman

**Author notes:** Correspondence to: Dr. Paul Hoffman, School of Philosophy, Psychology & Language Sciences, University of Edinburgh, 7 George Square, Edinburgh, EH8 9JZ, UK.

## Abstract

How does the brain code the meanings conveyed by language? Neuroimaging studies have investigated this by linking neural activity patterns during discourse comprehension to semantic models of language content. Here, we applied this approach to the production of discourse for the first time. Participants underwent fMRI while producing and listening to discourse on a range of topics. We used a distributional semantic model to quantify the similarity between different speech passages and identified where similarity in neural activity was predicted by semantic similarity. When people produced discourse, speech on similar topics elicited similar activation patterns in a widely distributed and bilateral brain network. This network was overlapping with, but more extensive than, the regions that showed similarity effects during comprehension. Critically, cross-task neural similarities between comprehension and production were also predicted by similarities in semantic content. This result suggests that discourse semantics engages a common neural code that is shared between comprehension and production. Effects of semantic similarity were bilateral in all three RSA analyses, even while univariate activation contrasts in the same data indicated left-lateralised BOLD responses. This indicates that right-hemisphere regions encode semantic properties even when they are not activated above baseline. We suggest that right-hemisphere regions play a supporting role in processing the meaning of discourse during both comprehension and production.

## Introduction

Both the production and comprehension of language are rooted in our conceptual knowledge of the world. At the neural level, this semantic processing is carried out through the activation of a widely distributed network of systems that store and retrieve knowledge (Binder & Desai, 2011; Lambon Ralph, Jefferies, Patterson, & Rogers, 2017; Pulvermüller, 2013). Traditionally, work on semantic representation has linked broad semantic categories (e.g., animals vs. tools) to regional engagement of semantic processing areas (Binder, Desai, Graves, & Conant, 2009; Martin, 2007). But more recently, the development of sophisticated multivariate pattern analysis (MVPA) techniques has allowed researchers to examine how conceptual knowledge is encoded across distributed patterns of neural activation (Kriegeskorte, Mur, & Bandettini, 2008; Norman, Polyn, Detre, & Haxby, 2006). These methods have moved the field from a coarse category-based approach to semantic representation into a fine-grained multidimensional understanding of the neural coding of meaning. They have also expanded the space of investigation from individual words and objects into the rich array of concepts and situations that occur in natural discourse.

In this study we use Representational Similarity Analysis (RSA), an MVPA technique that tests how similarity in the properties of stimuli can predict similarity in the neural activation patterns they elicit (Haxby, Connolly, & Guntupalli, 2014; Kriegeskorte et al., 2008). Previous studies have used RSA at the single-word level to investigate how semantic content is coded in the brain (Carota, Kriegeskorte, Nili, & Pulvermüller, 2017; Devereux, Clarke, Marouchos, & Tyler, 2013; Fischer-Baum, Bruggemann, Gallego, Li, & Tamez, 2017; Wang et al., 2018). The present study applies this approach to the discourse level and asks a critical but under-investigated question: is the coding of semantic content similar when people produce their own narratives, compared to when they listen to another person’s speech? In other words, does meaning in language comprehension and production rely on a common neural code?

MVPA has been used to investigate questions of semantic representation at different levels of stimulus complexity. Studies using stimuli at the single word (e.g., Carota et al., 2017; Devereux et al., 2013; Mitchell et al., 2008) and sentence level (e.g., Anderson et al., 2017; Pereira et al., 2018; Wang, Cherkassky, & Just, 2017) have demonstrated that patterns of neural activation can be predicted from vector-based semantic models. These include models that code conceptual relationships based on associated sensory-motor experiences (Binder et al., 2016; Fernandino et al., 2016) and models of natural language processing, which derive semantic representations from lexical statistics in large text corpora (Landauer & Dumais, 1997; Mikolov, Chen, Corrado, & Dean, 2013; Pennington, Socher, & Manning, 2014). Beyond the word and sentence levels, studies have used voxel-wise (de Heer, Huth, Griffiths, Gallant, & Theunissen, 2017; Huth, De Heer, Griffiths, Theunissen, & Gallant, 2016) and multivariate approaches to examine language comprehension at the discourse level. Such studies use more naturalistic stimuli, such as written stories (Dehghani et al., 2017; Wehbe et al., 2014), spoken narratives (Huth et al., 2016; Schrimpf et al., 2021; Zhang, Han, Worth, & Liu, 2020) or movies (Baldassano, Hasson, & Norman, 2018). These studies have revealed a widely distributed set of frontal, temporal, and parietal areas in which activation patterns can be predicted by the semantic content of language.

MVPA studies have shown that the neural representations of concepts are robust to changes in the stimuli used to evoke them. In other words, concepts elicit similar patterns of activation in the brain’s semantic network, irrespective of the stimuli used to evoke them. This cross-stimulus generalisation has been observed for object names vs. object pictures (Devereux et al., 2013; Fairhall & Caramazza, 2013); for written vs. spoken words (Liuzzi et al., 2017); for stories vs. movie stimuli (Baldassano et al., 2018); and for event descriptions that vary in lexical and syntactic content (Asyraff, Lemarchand, Tamm, & Hoffman, 2021). This is true even across languages – neural representations of concepts show cross-linguistic invariance at the individual word (Correia et al., 2014) and story-level (Dehghani et al., 2017). These findings are consistent with contemporary accounts of semantic cognition, which hold that higher-order conceptual knowledge is supported by “hub” regions that permit generalisation of knowledge across contexts, exemplars and modalities (Binder & Desai, 2011; Lambon Ralph et al., 2017; Margulies et al., 2016; Patterson, Nestor, & Rogers, 2007).

The critical question in our study is the degree to which neural semantic codes are similar across different language processing modes – specifically, comprehending versus producing speech. Theories of language production and comprehension suggest that these are interwoven processes that recruit overlapping neural networks (Hagoort, 2013; Kintsch & Vandijk, 1978; Levelt, Roelofs, & Meyer, 1999; Pickering & Garrod, 2013). At the level of discourse, both comprehension and production are thought to involve the construction of situation models which encode the entities and events described (Kintsch & Vandijk, 1978; Pickering & Garrod, 2004). In addition, one particular view postulates that comprehension uses a prediction-by-production mechanism, whereby the listener covertly imitates the linguistic form of the speaker’s utterances in order to derive their intentions, and then runs this through their production system to predict the speaker’s upcoming utterances (Pickering & Garrod, 2013). However, there is currently limited evidence that the neural representations engaged when people produce discourse are similar to those elicited when they listen to it. This is in part because, while MVPA studies of discourse comprehension are now commonplace, analogous studies of discourse production are rare.

Univariate neuroimaging studies suggest some differences in the neural networks activated during comprehension and production. While production and comprehension of speech activate common language areas in the left hemisphere, narrative comprehension tends to activate right frontal and temporal regions more than production (AbdulSabur et al., 2014; Awad, Warren, Scott, Turkheimer, & Wise, 2007; Silbert, Honey, Simony, Poeppel, & Hasson, 2014). These findings align with theoretical views that while speech comprehension engages a bilateral network, language production is more left-lateralised (Federmeier, 2007; Hickok & Poeppel, 2007; Lambon Ralph, McClelland, Patterson, Galton, & Hodges, 2001; Poeppel, Emmorey, Hickok, & Pylkkänen, 2012; Schapiro, McClelland, Welbourne, Rogers, & Lambon Ralph, 2013). More recently, a series of studies have investigated correlations in neural activity between a speaker producing a story and a listener comprehending it (Jiang et al., 2012; Liu et al., 2017; Silbert et al., 2014; Stephens, Silbert, & Hasson, 2010). These indicate a more bilateral picture, with significant production-comprehension temporal coupling in left and right prefrontal, temporal and inferior parietal cortices. This inter-subject coupling suggests a large degree of shared neural processing across comprehension and production. However, these studies have investigated correlations between different individuals and not within the same individual performing different language tasks, which is important given evidence for widespread individual differences in language neuroanatomy (Fedorenko, Hsieh, Nieto-Castañón, Whitfield-Gabrieli, & Kanwisher, 2010). In addition, inter-subject correlations in neural activity could be due to shared processes at any level of the language system and do not indicate where commonalities are due specifically to its semantic content. Here, we used RSA to investigate the degree to which neural patterns during speech production and comprehension align with a single model of semantic content. In the scanner, participants listened to pre-recorded speech samples, and produced extended passages of speech on a variety of topics. We used a distributional semantic model (latent semantic analysis; Landauer & Dumais, 1997) to quantify the degree to which different speech chunks contained similar semantic content. We then used RSA to assess how well semantic similarity could predict similarity in the neural responses evoked during different periods of heard or spoken language. Critically, as well as performing these analyses separately for comprehension and production data, we conducted a cross-task analysis that looked at similarities between comprehension and production periods (within participants). In this way, we were able to identify brain regions which share a common neural code for semantic content during speaking and listening.

We used a searchlight approach to seek semantic-coding regions across the whole brain. In addition, we investigated effects in a set of regions of interest known to be involved in semantic processing. These comprised parts of the anterior temporal lobes, which act as a hub for representation of semantic knowledge (Lambon Ralph et al., 2017; Patterson et al., 2007), inferior prefrontal and posterior temporal regions that regulate access to this knowledge (Badre & Wagner, 2007; Jefferies, 2013) and the angular gyrus, a key node in the default mode network which is involved in constructing mental models of events and ongoing experiences (Binder & Desai, 2011; Yeshurun, Nguyen, & Hasson, 2021). By investigating effects in the left and right hemisphere homologues of these regions, we were also able to probe the extent of lateralization during production and comprehension of speech.

## Method

### Participants

25 young adults participated in the study in exchange for payment. Their mean age was 24 (SD = 4.4, range = 18-35), all were native English speakers and they were all classified as right-handed using the Edinburgh Handedness Inventory (Oldfield, 1971). The study was approved by the Psychology Research Ethics Committee of the University of Edinburgh and all participants gave informed consent. Unrelated analyses of this dataset, investigating effects of discourse coherence and other psycholinguistic properties, have been reported previously (Morales, Patel, Tamm, Pickering, & Hoffman, 2022; Wu, Morales, Patel, Pickering, & Hoffman, 2022). In contrast, the present study used multivariate analyses to investigate the neural representation of discourse content.

### Materials

In the speaking task, 12 prompts were used to elicit discourse from participants. Each prompt probed common semantic knowledge on a specific topic (e.g., *Describe how you would make a cup of tea or coffee*; see Supplementary Materials for a complete list of prompts in both tasks). This discourse speaking condition was contrasted with a baseline of automatic speech that involved repeated recitation of the English nursery rhyme, Humpty Dumpty (AbdulSabur et al., 2014; Blank, Scott, Murphy, Warburton, & Wise, 2002; Hoffman, 2019). For the listening task, a different set of 12 topics was used. We generated audio samples of discourse using transcripts of speech provided by participants in a previous behavioural study, in response to 12 topic prompts (Hoffman, Loginova, & Russell, 2018a). For each topic, we selected two different responses provided by different participants in the Hoffman et al. (2018) study. One sample was highly consistent with the topic and the other deviated somewhat from the specified topic, as indicated by global coherence measured using the methods of Hoffman et al (Hoffman, 2019; Hoffman et al., 2018a). We did this to increase the variability in the semantic content of the stimuli. We divided the listening samples into two counterbalanced sets, so that each participant would hear one of the two sets. To generate the audio recordings that would be presented in the scanner, the 24 transcripts were read aloud by the same male native English speaker and edited so that their duration was 50s each (a sample transcript is provided in Supplementary Materials). A 10s recording of the same speaker reciting the Humpty Dumpty rhyme was also made to provide a baseline listening condition.

### Design and procedure

Each participant completed two speaking and two listening runs, presented in an alternating sequence. The order of runs was counterbalanced over participants. Each run lasted approximately 8 minutes and included six discourse trials and five baseline trials, with the order of discourse trials in each run fully randomised for each participant.

Speaking trials began with the presentation of a written topic prompt on screen for 6s (see Figure 1A). Participants were asked to prepare to speak during this period and to start speaking when a green circle replaced the prompt in the centre of screen. They were instructed to speak about the topic for 50s, after which a red cross would replace the green circle. At this point participants were instructed to wait for the next prompt to appear on screen. The procedure for listening trials was the same, except participants were asked to listen to the speaker attentively for 50s while the green circle was on screen. For the baseline automatic speech conditions, participants were instructed to recite or to listen to the Humpty Dumpty rhyme for a 10s period. When speaking, they were asked to start reciting again from the beginning if they reached the end of the nursery rhyme before the 10s had elapsed. The baseline conditions therefore involved grammatically well-formed continuous speech, but without the requirement to generate or understand novel discourse. All trials were presented with an inter-stimulus interval jittered between 3s and 7s (M = 5s).

**Figure 1.**
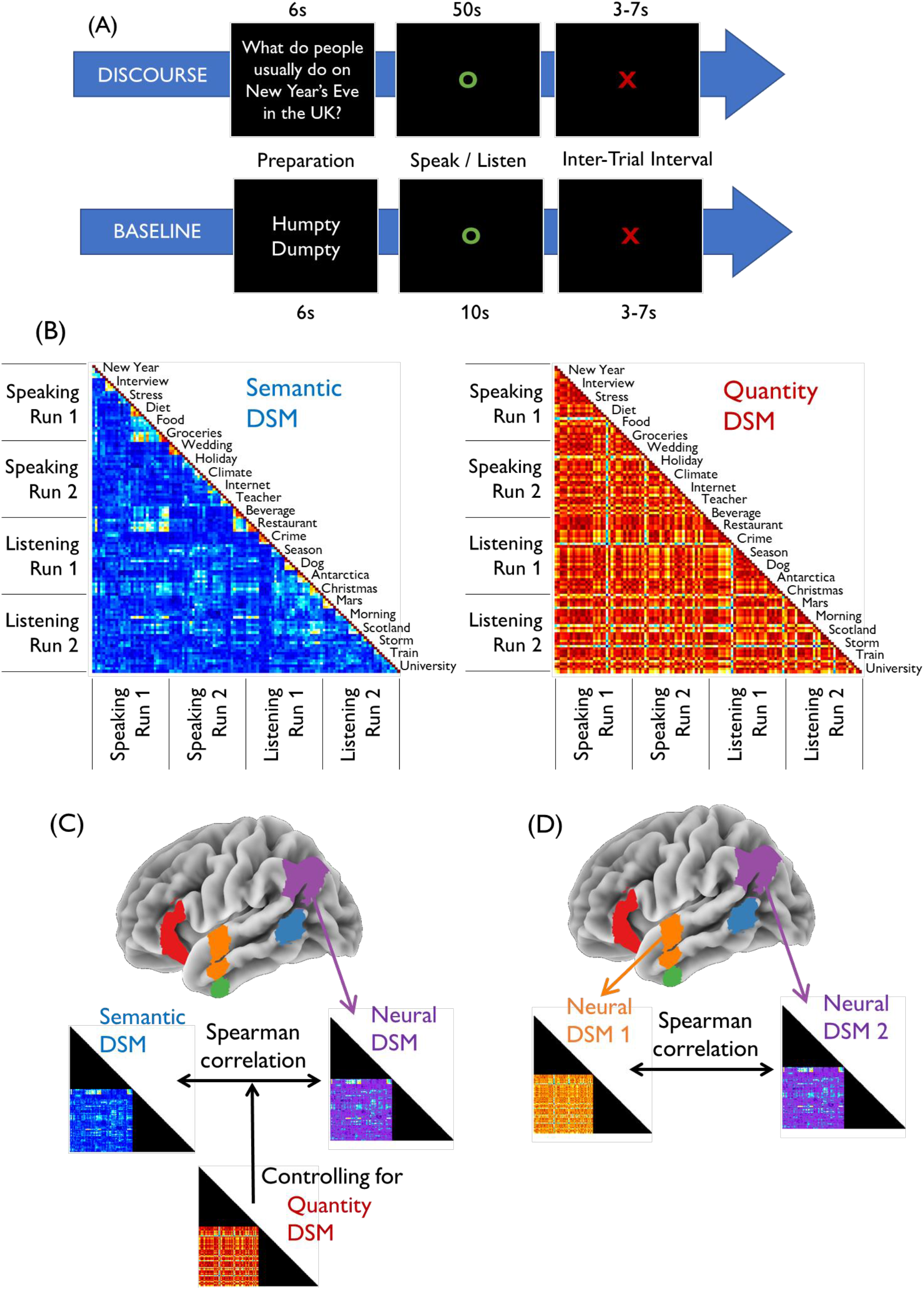
(A) Structure of trials in the experiment. On each discourse trial, participants were presented with a topic prompt for 6s and were then required to either speak about this topic for 50s or listen to a recording of another person speaking about it for 50s. In the baseline condition, participants either recited or listened to a well-known British nursery rhyme. (B) Examples of semantic and quantity dissimilarity matrices. Topic labels indicate different discourse topics, each of which was divided into five 10s segments. The semantic DSM codes how similar speech segments are in lexical-semantic content. The quantity DSM (used as a control) codes how similar segments are terms of the number of words they contain. (C) Procedure for main RSA analyses. The analysis tested the relationship between a neural DSM, generated by comparing local activation patterns for different speech segments, and the semantic DSM, while controlling for the effects of the quantity DSM. (D) Procedure for comparison of neural DSMs. This analysis tests the relationship between two neural DSMs obtained from different brain regions. Speak1 = speech production run 1; Speak2 = speech production run 2; Listen1 = comprehension run 1; Listen2 = comprehension run 2. DSM = dis-similiarity matrix.

**Figure 2:**
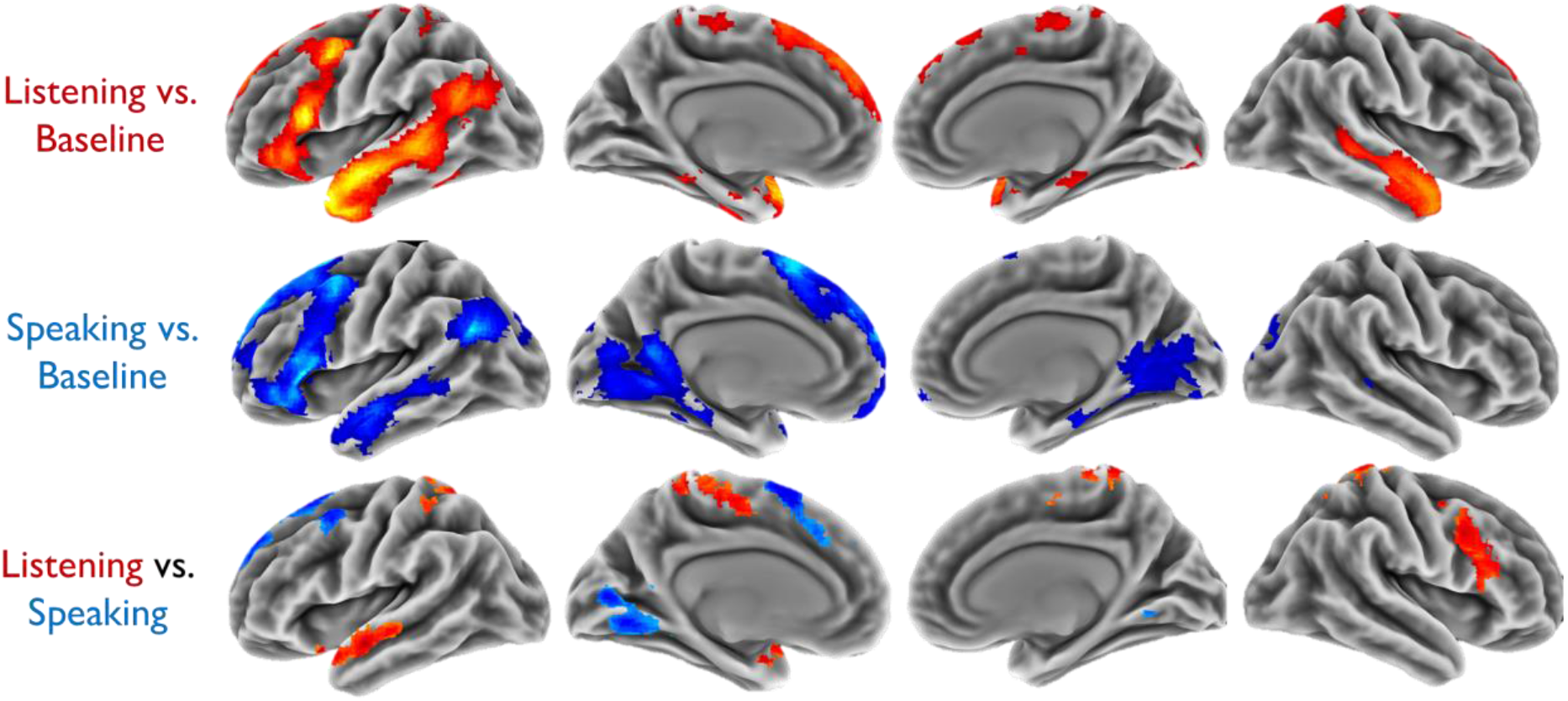
Univariate activation contrasts. Areas activated by each discourse task compared with its baseline, and for the direct contrast of speaking vs. listening. Results are shown at cluster-corrected p < 0.05. Peak activation co-ordinates are reported in Supplementary Table 1.

Before scanning, participants were presented with training trials to familiarise them with the tasks. To ensure attention during listening runs, they were told that they would receive a memory test on the material after scanning. In this test, participants answered 12 multiple choice questions, one for each topic presented during listening. They were also asked to rate how well they could hear the speech samples in the scanner from 1 (inaudible) to 7 (perfectly audible).

### Processing of speech samples

Spoken responses were digitally recorded with an MRI-compatible microphone and processed with noise cancellation software (Cusack, Cumming, Bor, Norris, & Lyzenga, 2005) to reduce noise from the scanner. They were then manually transcribed. For analysis, each 50s response was divided into 5 segments of 10s duration.

For each participant, we constructed two dissimilarity matrices (DSMs) based on the properties of the speech produced or heard in each 10s segment. Our main DSM indexed the semantic relatedness of speech segments using Latent Semantic Analysis (LSA) (Landauer & Dumais, 1997). Like other distributional models of semantic representation (Mikolov et al., 2013; Pennington et al., 2014), LSA represents words as embeddings in a high-dimensional semantic space. The proximity of two words in this space indicates the degree to which they are used in similar contexts in natural language, which is taken as a proxy for similarity in meaning. We used LSA vectors generated from the British National Corpus, which we have used previously for analyses of discourse production (Hoffman, 2019; Hoffman et al., 2018a). As in previous studies, a vector representation for each speech segment was generated by averaging the LSA vectors of all of the words contained in the segment (excluding function words and weighting vectors by their log frequency in the segment and their entropy in the original corpus; Hoffman et al., 2018a). Once a vector had been calculated for each speech segment, a semantic DSM was calculated using a cosine similarity metric. An example of a semantic DSM for a single participant is shown in Figure 1B and DSMs for all participants are provided in Supplementary Figure 1. Each participant’s semantic DSM was unique as each participant produced different information in response to the topic prompts.

We computed a second DSM that captured variation in the quantity of speech produced or heard in different segments. Here, dissimilarity between two segments was defined as the difference in the number of words contained in the two segments; thus, segments containing a similar quantity of speech were represented as similar to one another. A quantity DSM for a single participant is shown in Figure 1B. We included the quantity DSM as a covariate in our semantic analyses, to ensure that our effects were being driven by the semantic content of the discourse and were not confounded by variation in the amount of speech participants were processing.

### Image acquisition and processing

Participants were scanned on a 3T Siemens Prisma scanner using a 32-channel head coil. fMRI data were acquired at three echo times (13ms, 31ms, and 48ms) using a whole-brain multi-echo acquisition protocol. Data from these three echo series were weighted and combined, and the resulting time-series were denoised using independent components analysis (ICA). This approach improves the signal quality in regions that typically suffer from susceptibility artefacts (e.g., the ventral anterior temporal lobes) and is helpful in reducing motion-related artefacts (Kundu et al., 2017). The TR was 1.7 s and images consisted of 46 slices with an 80 x 80 matrix and isotropic voxel size of 3mm. Multiband acceleration with a factor of 2 was used and the flip angle was 73°. Four runs of 281 volumes (477.7s) were acquired. A high-resolution T1-weighted structural image was also acquired for each participant using an MP-RAGE sequence with 1mm isotropic voxels, TR = 2.5 s, TE = 4.6 ms.

Images were pre-processed and analysed using SPM12 and the TE-Dependent Analysis Toolbox 0.0.7 (Tedana) (DuPre et al., 2019). Estimates of head motion were obtained using the first BOLD echo series. Slice-timing correction was carried out and images were then realigned using the previously obtained motion estimates. Tedana was used to combine the three echo series into a single-time series and to divide the data into components classified as either BOLD-signal or noise-related based on their patterns of signal decay over increasing TEs (Kundu et al., 2017). Components classified as noise were discarded. After that, images were unwarped with a B0 fieldmap to correct for irregularities in the scanner’s magnetic field. Finally, functional images were spatially normalised to MNI space using SPM’s DARTEL tool (Ashburner, 2007).

For univariate analysis, images were smoothed with a kernel of 8mm FWHM. Data were treated with a high-pass filter with a cut-off of 128s and the four experimental runs were analysed using a single general linear model. Four types of speech block were modelled: discourse speaking (50s), baseline speaking (10s), discourse listening (50s), and baseline listening (10s). Prompt presentation periods were modelled as blocks of 6s, with separate regressors for discourse prompts and Humpty Dumpty prompts. Covariates consisted of six motion parameters and their first-order derivatives.

For multivariate analysis, images were smoothed with a kernel of 4mm FWHM, as a small amount of smoothing has been shown to improve the sensitivity of multivariate analyses (Gardumi et al., 2016; Hendriks, Daniels, Pegado, & Op de Beeck, 2017). Each run was modelled in a separate general linear model. Regressors for prompts and baseline conditions were the same as in the univariate analysis. Discourse blocks were divided into segments of 10s duration (5 per prompt) and each segment was modelled using a separate regressor. Thus, we obtained a separate beta map for each 10s segment of speech heard/produced by each participant. Baseline activation was not subtracted from these results, so patterns represented changes in BOLD signal relative to the model intercept (rest). The beta maps were converted into *t*-maps for RSA analysis. Motion covariates were included as in the univariate analysis.

### Regions of interest

In addition to whole-brain analyses, we defined five anatomical regions in the left and right hemispheres, shown in Figure 3A. We chose these regions because they have all been reliably implicated in semantic cognition (Binder & Desai, 2011; Lambon Ralph et al., 2017). Four of the five ROIs were defined using probability distribution maps from the Harvard-Oxford brain atlas (Makris et al., 2006), including all voxels with a >30% probability of falling within the following regions:

1. Inferior frontal gyrus (IFG): the pars orbitalis and pars triangularis regions of inferior frontal gyrus, with voxels more medial than x = ±30 removed to exclude medial orbitofrontal cortex
2. Lateral anterior temporal lobe (lATL): the anterior division of the superior and middle temporal gyri
3. Ventral anterior temporal lobe (vATL): the anterior division of the inferior temporal and fusiform gyri
4. Posterior middle temporal gyrus (pMTG): the temporo-occipital part of the middle temporal gyrus

**Figure 3:**
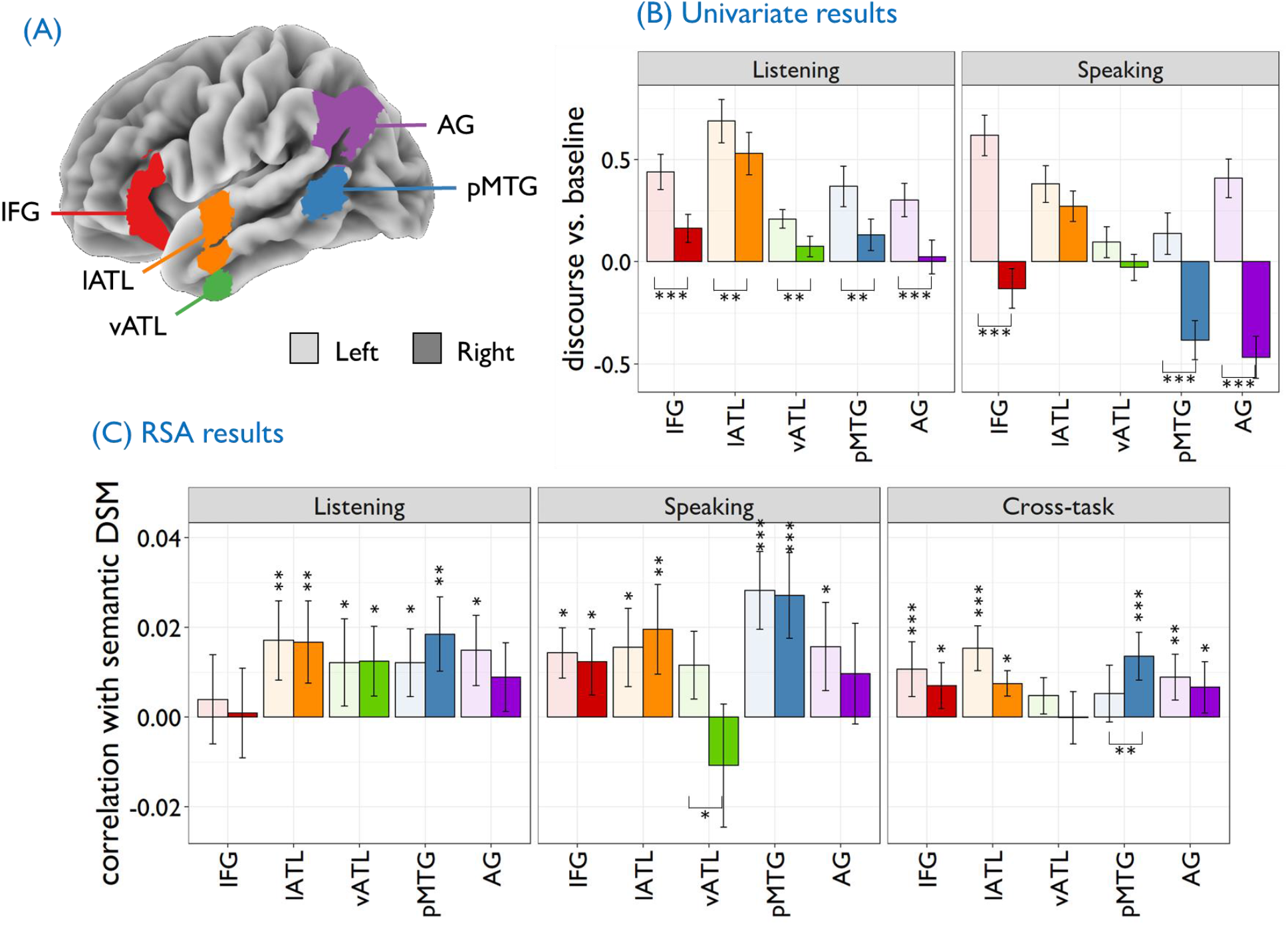
Region of interest analyses. (A) ROI locations shown on the left hemisphere. Regions were selected based on known involvement in semantic processing and defined using an anatomical atlas (see Method for details). (B) Univariate activation values for discourse vs. baseline, where pale bars represent left-hemisphere activation and bright bars represent the right hemisphere. Asterisks below the x-axis indicate significant hemispheric differences. (C) RSA results, showing the Fisher-transformed correlation between the semantic DSM and neural DSM in each ROI. Asterisks above the x-axis indicate correlations significantly greater than zero, while those below the x-axis indicate significant hemispheric differences. * = p < 0.05; ** = p < 0.01; *** = p < 0.001, all FDR-corrected. Error bars show 1 SEM. IFG = inferior frontal gyrus; lATL = lateral anterior temporal lobe; vATL = ventral anterior temporal lobe; pMTG = posterior middle temporal gyrus; AG = angular gyrus.

The final ROI covered the angular gyrus (AG) and included voxels with a >30% probability of falling within this region in the LPBA40 atlas (Shattuck et al., 2008). A different atlas was used in this case because the AG region defined in the Harvard-Oxford atlas is small and does not include parts of the inferior parietal cortex typically implicated in semantic processing.

### Univariate analyses

We obtained whole-brain activation maps for speech production (discourse minus baseline) and speech comprehension (discourse minus baseline). We computed a subtraction of these two maps to identify differences between speaking and listening at the discourse level. For these analyses we employed a voxel-height threshold of *p* < 0.005 for one-sample *t*-tests with correction for multiple comparisons at the cluster level, using SPM’s random field theory. The direct comparison of speaking and listening was restricted to voxels that showed an effect of discourse > baseline in at least one of the two tasks (at a liberal threshold of *p* < 0.05 uncorrected). In addition to whole-brain analysis, contrast estimates for discourse minus baseline were extracted for each ROI in each task and were entered into a 2 x 2 x 5 (task x hemisphere x ROI) ANOVA.

### Representational similarity analysis

RSA was used to investigate the degree to which neural patterns were predicted by the semantics of the discourse that participants heard and produced. Analyses were performed using CoSMoMVPA (Oosterhof, Connolly, & Haxby, 2016). Searchlight analyses were run using a spherical searchlight of radius 12mm (4 voxels) and ROI analyses were performed in the 10 semantic regions described previously. Because the ROIs varied substantially in size, a voxel selection criterion was applied to ensure that all ROI analyses used the same number of voxels. For each participant, voxels in each ROI were ordered by their effect size in the contrast of discourse over baseline and the 100 most active voxels in each ROI were selected for the RSAs. This ensured that all ROI analyses were equally powered and were based on the voxels that were most engaged by the task.

Separate RSAs were performed for the listening and speaking tasks, as well as a cross-task analysis that tested pattern similarity across comprehension and production. For listening, a neural DSM was generated by calculating 1 minus Pearson correlations between activation patterns for pairs of speech segments presented during the listening task. To avoid any confounding effects of temporal auto-correlation in the BOLD signal, only pairs of segments from different scanning runs were compared (Mumford, Davis, & Poldrack, 2014) (see Figure 4A). This also meant that segments from within the same speech topic were never compared. Activation patterns were mean-centred within each run, ensuring that each voxel had a mean activation of zero (as recommended by Diedrichsen & Kriegeskorte, 2017). For the speaking task, the same process was followed but pairs were taken from the speaking runs (see Figure 4B). For the cross-task analysis, each pair consisted of one speaking segment and one listening segment (see Figure 4C). Thus, the cross-task analysis tested the degree to which similarity in neural patterns across language tasks could be explained by their semantic content.

**Figure 4:**
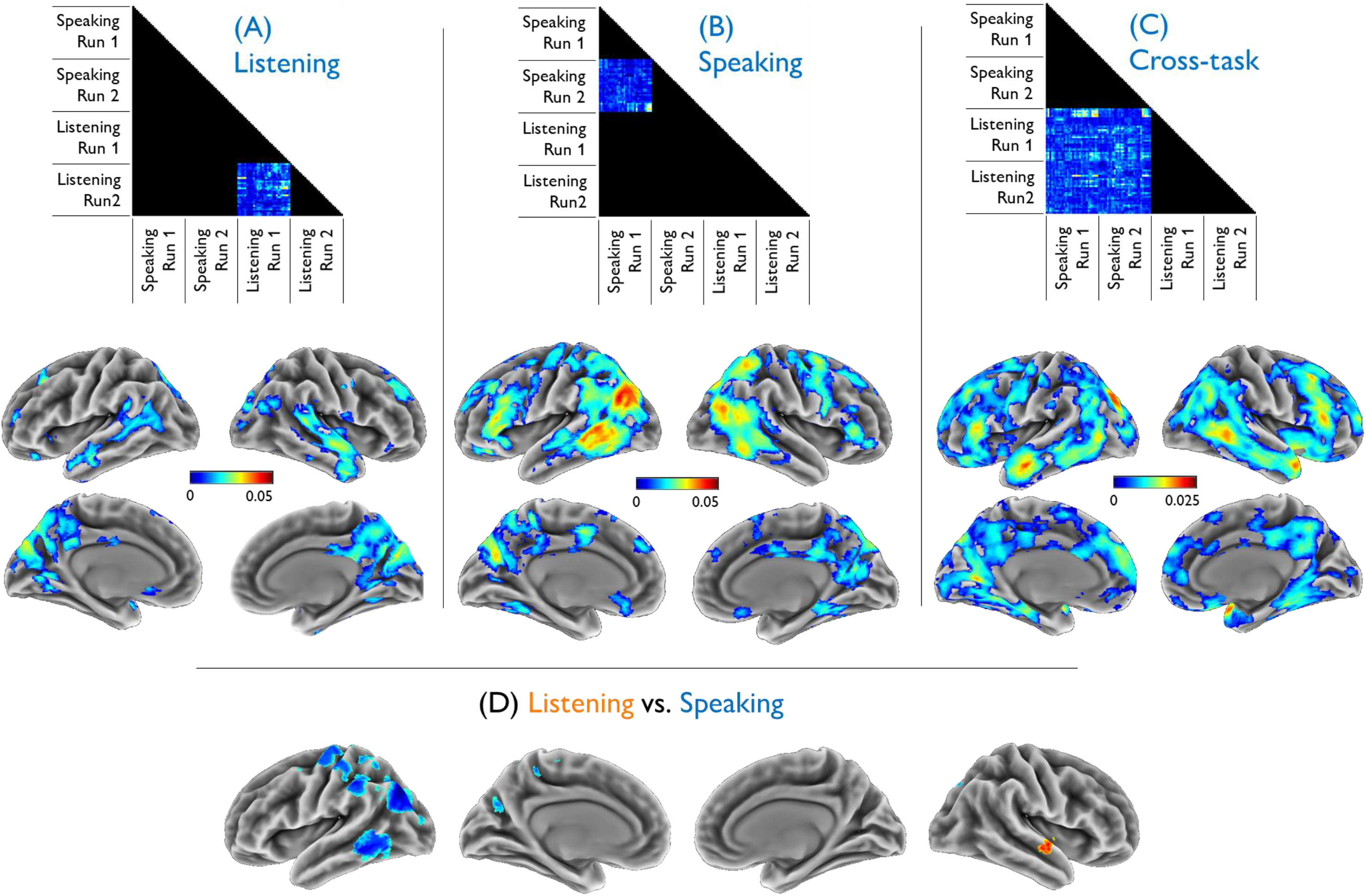
Searchlight analyses. (A-C) The top panel in the figure indicates which parts of the DSMs were used in each analysis. DSMs consist of pairwise comparisons between different 10-second segments of discourse processing. In the Listening and Speaking analyses, we used pairs taken from the same language task but taken from different scanning runs (since segments in the same run may be affected by temporal autocorrelation in the BOLD signal; see Methods). In the cross-task analysis, pairs consisted of one Speaking and one Listening segment (which were always from different scanning runs). Brain maps show regions where the correlation between neural and semantic DSMs was significantly greater than zero (at cluster-corrected p < 0.05). Colour scales show the group-average Fisher-transformed correlation coefficient between semantic and neural DSMs; note that the scale is different in the cross-task analysis. Peak effect co-ordinates are reported in Supplementary Table 2. (D) The bottom panel shows differences in correlation coefficients between Listening and Speaking analyses, where differences were statistically significant (at cluster-corrected p < 0.05).

The association between neural DSMs and their corresponding semantic DSMs was measured using partial Spearman correlations, which were Fisher-transformed prior to statistical inference. Analyses included the quantity DSMs as a covariate, ensuring that effects were not dependent on the number of words in each segment (see Figure 1C). To test the degree to which our results were specific to semantic processing, control analyses were conducted using quantity DSMs as the predictor and semantic DSMs as the covariate. The results of these are reported in Supplementary Materials.

To determine whether semantic DSMs significantly predicted neural dissimilarity patterns, permutation tests were performed using the two-stage method introduced by Stelzer et al. (2013). For each participant, we calculated Spearman partial correlations using a semantic DSM in which the order of segments had been randomly permuted within each run. This process was repeated 100 times to provide a distribution of results for each participant under the null hypothesis. Following this, a Monte Carlo approach was taken to generate a null distribution at the group level. We randomly selected one correlation map from each participant’s null distribution and averaged these to give a group mean. This process was repeated 10,000 times to generate a distribution of the expected group correlations under the null hypothesis. In ROI analyses, the position of the observed result in this null distribution was used to determine a p-value (e.g., if the observed accuracy was greater than 99% of values in the null distribution, this would represent a p-value of 0.01). For searchlight analyses, observed and null maps were entered into CoSMoMVPA’s Monte Carlo cluster statistics function, which returned a statistical map corrected for multiple comparisons using threshold-free cluster enhancement (Smith & Nichols, 2009). These maps were thresholded at corrected *p* < 0.05. Finally, to test whether any regions showed differences in the strength of the semantic correlations between tasks, we calculated a difference map by subtracting the correlations in the speaking task from those in the listening task. We then tested for regions where the difference was significantly greater or less than 0, using the permutation methods described above.

### Assessing lateralisation in activation and RSA maps

To formally assess the similarity of effects in the left and right hemispheres, we measured spatial correspondence between significant voxels in each hemsiphere. To do this, we first binarised each thresholded activation/RSA image so that voxels showing a significant effect had a value of 1 and other voxels a value of 0. We then created a mirror image of each binary map by flipping it on the x-axis (for similar approaches, see Alam, Karapanagiotidis, Smallwood, & Jefferies, 2019; Hoffman & Morcom, 2018). We calculated the Jaccard similarity between the original map and its flipped version. Jaccard similarity quantifies the degree to which two analyses share the same significant voxels, with 1 indicating that significant voxels in both analyses are identical and 0 indicating no voxels in common (Maitra, 2010). When applied to our images, this gave a measure of the degree of spatial overlap in the effects in each hemisphere. High values indicate highly symmetrical effects, while low values indicate greater divergence between hemispheres. We obtained these measures for each of our activation contrasts and RSA searchlight analyses.

### Comparison of similarity structure in different brain regions

Finally, we investigated the relationship between neural DSMs in different ROIs (see Figure 1D). The purpose of this analysis was to explore the degree to which different parts of the semantic network represented discourse content in a similar way. For each participant, Spearman correlations were calculated between pairs of neural DSMs, for all possible pairs of ROIs. This resulted in a correlation matrix that coded the degree to which pairs of ROIs shared similar neural DSMs. These were then averaged over participants to give a single correlation matrix for the whole group. This process was performed within the listening and speaking tasks, as well as across tasks. To visualise the relationships between ROIs, group correlation matrices were converted to distance matrices and hierarchical cluster analyses were performed using R, with Ward’s distance metric.

## Results

We performed a series of analyses on fMRI data in which participants produced and listened to speech samples on a range of topics (see Figure 1). After reporting basic characteristics of the language samples, we report the results of univariate analyses which investigated the degree to which brain regions were engaged during discourse comprehension and production, compared with a baseline of automatic speech. Following this, we use RSA to identify regions in which similarities in activation patterns are predicted by similarities in the semantic content of language. As well as performing these analyses separately on listening and speaking data, we perform a cross-task analysis that tests for semantic neural coding that generalises across language tasks. Finally, we directly compare neural similarity patterns across semantic-related brain regions to investigate the degree to which different regions are similarly influenced by discourse content.

### Characteristics of speech

In the discourse production task, participants spoke about a series of topics for 50s at a time (see Method for details). They produced a mean of 124 words per topic (SD = 25, range = 67-196; for distribution over participants, see Supplementary Figure 2). In the discourse comprehension task, participants listened to recordings of another person speaking about a different set of topics. These recordings contained a mean of 141 words per topic (SD = 20, range = 102-170). Example speech samples from comprehension and production tasks are provided in Supplementary Materials. After scanning, participants received 12 comprehension questions relating to the speech presented in the listening task. They answered 10/12 questions correctly on average. They also provided audibility rating with a mean rating of 5.5/7, suggesting that they were able to understand the discourse samples presented in the scanner.

### Univariate analyses

Whole-brain activation maps are shown in Figure 2. These show effects for each discourse task relative to a “low-level” speech baseline (reciting/listening to a familiar nursery rhyme). The baseline conditions involved language processing and so had similar perceptual and motor demands to the discourse tasks, but they did not require participants to process novel, meaningful verbal information. In line with previous studies, the results indicate close correspondence in the areas recruited for production and comprehension of novel discourse, particularly in the left hemisphere. Both tasks activated similar left-lateralised networks associated with semantic processing, including lateral and medial prefrontal cortex, lateral temporal cortex, the ventral anterior temporal lobe and the angular gyrus. Listening produced significantly greater activation than speaking in right prefrontal regions, left superior temporal sulcus and bilateral post-central gyrus. Speech production was associated with greater activity in the cerebellum, medial prefrontal cortex and the occipital lobe. The overall picture, however, was that participants engaged broadly similar networks whether they were speaking or listening.

We also assessed activation in five regions of interest (ROIs) that are frequently implicated in semantic processing, and their right-hemisphere homologues (see Figure 3A). Results are shown in Figure 3B and reveal a left-lateralised pattern for both tasks. 2 x 5 x 2 ANOVA showed a main effect of hemisphere (*F*(1,24) = 64.4, *p* < .001; left > right) as well as ROI (*F*(4,96) = 20.7, *p* < .001), and task (*F*(1,24) = 5.7, *p* = .026; listening > speaking). All of the interactions between these factors were also significant (*F* > 6.9, *p* < .001). FDR-corrected post-hoc tests indicated that activation was left-lateralised for both tasks in most regions (see Figure 3B). In the IFG, pMTG and AG, this effect was larger in the speaking task, with right-hemisphere regions deactivating during production, relative to the baseline speech condition. Similar results were obtained when discourse was contrasted against rest rather than baseline speech conditions (see Supplementary Figure 3). Thus, univariate analyses suggest that the left hemisphere is dominant in semantic aspects of discourse processing, particularly when participants produced, rather than heard, discourse.

### RSA searchlights

These analyses investigated whether similarity in the activation produced during different passages of speech can be explained by similarity in the semantic content of those passages. The RSA method assesses this by first computing a neural dis-similarity matrix (DSM) that measures the dis-similarity (1 – Pearson correlation) between activation patterns elicited at different points during task performance. To generate neural DSMs, we divided each 50s discourse period into 5 segments of 10s duration. We calculated the pairwise dis-similarities between the activation patterns evoked during these segments of speech (see Methods for details). We then tried to predict the values in the neural DSM using a semantic DSM, which coded dis-similarity between the content of segment using an established vector-based model of semantics (Landauer & Dumais, 1997). We also controlled for similarity in the quantity of speech contained in each segment (see Figure 1C and Methods). This process was repeated across a series of “searchlights” to build a whole-brain map of semantic-neural correlations. As shown in Figure 4, we performed three separate analyses: comparing listening segments with other listening segments, comparing speaking segments with other speaking segments and, in a cross-task analysis, comparing speaking segments with listening segments. Results of the three searchlight analyses are shown in Figure 4. When participants listened to discourse, semantic similarity predicted neural similarity in regions along the length of the superior temporal sulci bilaterally. In the anterior temporal lobes, effects extended ventrally into the middle and inferior temporal gyri, particularly in the right hemisphere. Significant correlations were also observed in a large area of posterior medial cortex encompassing posterior cingulate, cuneus and precuneus. These results converge with those of previous studies in indicating that the bilateral lateral temporal lobe regions, as well as other default mode network regions (posterior cingulate, medial prefrontal cortex), track semantic content during spoken narrative comprehension (Schrimpf et al., 2021; Zhang et al., 2020).

The second analysis extended the use of semantic RSA to the domain of discourse production for the first time. Here, activity in a more extensive set of regions was found to correlate with semantic similarity (Figure 4B). The strongest correlations were found in left pMTG and inferior parietal cortex, with significant associations also found in IFG, the default mode regions of posterior cingulate and ventromedial prefrontal cortex, as well as parts of the intraparietal sulci and motor cortices. Thus, when individuals produce discourse on similar topics, this is reflected in similar activation patterns within a widely distributed brain network. As in the listening analysis, the distribution of significant regions was largely bilateral, in contrast to the highly left-lateralised pattern observed in the univariate analysis (this effect is quantified in the next section).

Finally, the cross-task analysis tested for predictive effects of semantic similarity when comparing activation patterns across comprehension and production tasks. Significant effects here indicate that similarities in activation across language tasks can be predicted by the underlying semantic content of the discourse being processed. We interpret this as evidence for common coding of language content across speaking and listening tasks. The cross-task analysis revealed semantic-neural correlations in an extensive and bilateral set of regions (Figure 4C). These include regions classically associated with semantic and language processing, such as the anterior and posterior temporal lobes and lateral prefrontal cortices, but also nodes of the default mode network (inferior parietal lobes, posterior cingulate and medial prefrontal cortex) and left hippocampus and parahippocampal gyrus. There are two key implications of these results. First, they indicate that an array of brain regions encodes the content of one’s own speech in a similar way to the content of speech produced by others, suggesting that the semantics of comprehension and production share a common neural code. Second, they indicate that activation in right-hemisphere regions is as sensitive as left-hemisphere regions to the meaning of discourse. Thus, while univariate analyses suggest that the right hemisphere is less selectively engaged during discourse processing, its activation is nevertheless predicted by the semantic content, suggesting a potential functional role for these regions.

Although the cross-task analysis identified many of the same regions as the separate task-specific analyses, more voxels exceeded the significance threshold in this analysis. This is probably because a greater number of speech segment pairs were available when comparing across different tasks (as shown in the top panel of Figure 4). This allowed weaker semantic-neural correlations to reach statistical significance in the cross-task analysis. It is important to note that, although more voxels showed a statistically significant effect, the magnitudes of the semantic-neural correlations in the cross-task analysis were generally lower than in the separate listening and speaking analyses.

Figure 4D shows regions where RSA effects differed significantly between the listening and speaking analyses. Speaking produced significantly stronger correlations with semantic DSMs in parts of left posterior temporal and inferior parietal cortex. Listening produced stronger correlations in a small portion of right anterior STG. However, most of the regions identified also showed a cross-task semantic effect, suggesting they code discourse content in a similar way across speaking and listening. Differences in the correlation strength between tasks may indicate differences in how strongly these shared representations are engaged by different language tasks.

In the control analyses, neural DSMs were predicted used the quantity DSMs (which class speech segments as dis-similar if they contain very different numbers of words) rather than semantic DSMs. In the listening task, effects were found bilaterally in ventral premotor and superior temporal regions, centred on Heschl’s gyrus (see Supplementary Figure 4). The cross-task analyses revealed similar small clusters in the left and right superior temporal lobes, again centred on primary auditory cortex. Analysis of the speaking task revealed a more distributed set of regions, including bilateral motor and premotor cortices, some medial parts of the default mode network and left inferior parietal cortex, though with weaker correlations than those seen in the main analyses. None of these analyses revealed the patterns of temporal, inferior prefrontal and parietal effects seen in the main analysis. Thus, the control analyses suggest that effects in Figure 4 are specific to predictors that capture high-level conceptual content of language and not its lower-level properties.

### Assessing lateralisation in activation and RSA maps

Figure 5 shows left-right Jaccard similarity metrics for each analysis. These values quantify the degree to which the pattern of significant effects was symmetrical across hemispheres. A value of 1 would indicate that for every significant voxel in the left hemisphere, its equivalent voxel in the right hemisphere was also significant (and vice-versa). Conversely, 0 would indicate that whenever a left-hemisphere voxel was significant, its right-hemisphere equivalent was not significant (and vice-versa). Similarity values were substantially higher for the RSA effects than the activation contrasts, particularly in the Speaking and Cross-task analyses. This supports our assertion that RSA analyses reveal a more bilateral pattern of involvement in discourse processing, which is different to the left-lateralised effects seen in univariate contrasts.

**Figure 5:**
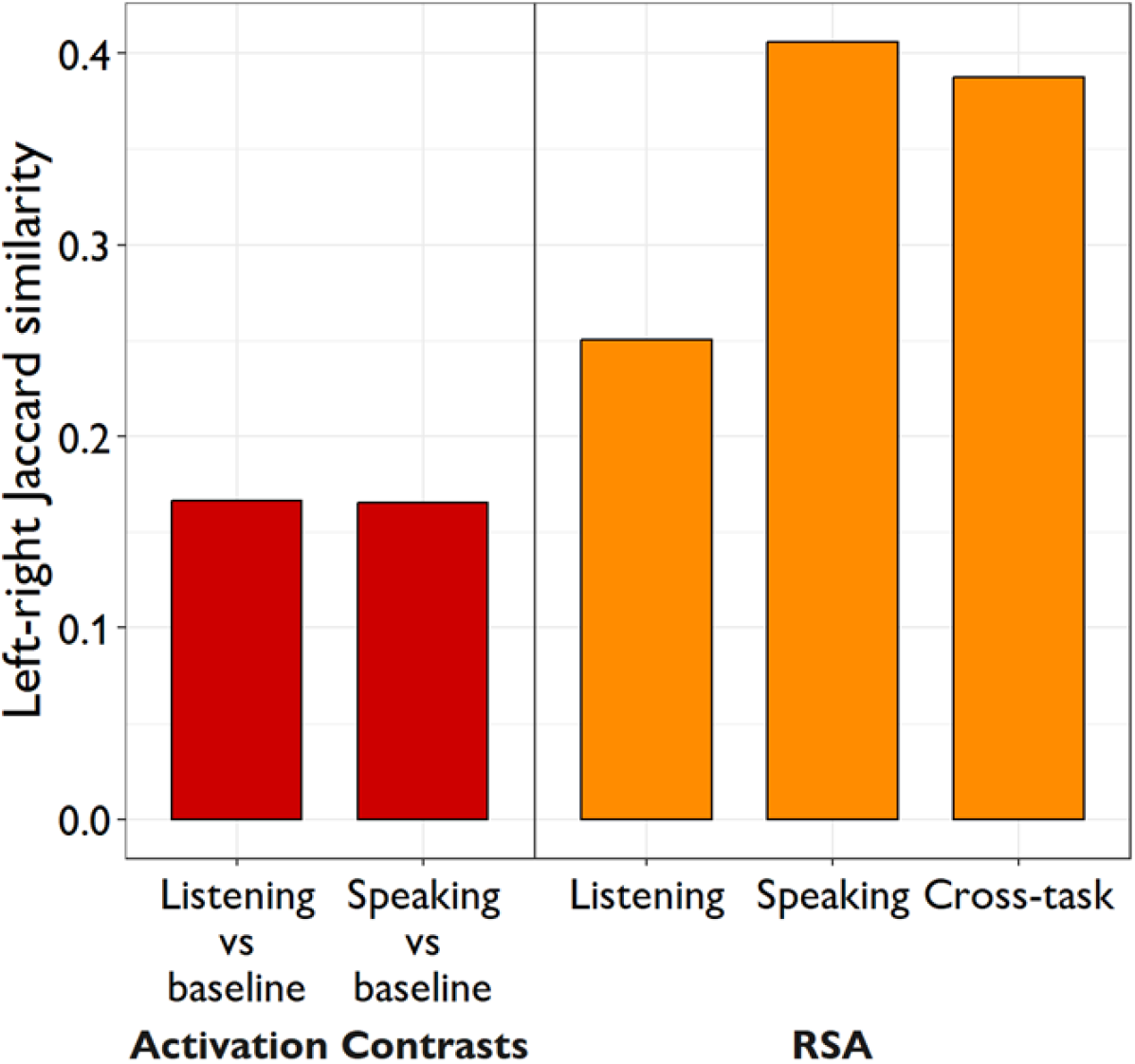
Similarity in the spatial distribution of effects in left and right hemispheres. Jaccard similarity values in this plot indicate the degree to which the same regions showed significant effects in the left and right hemispheres. The higher the Jaccard similarity, the more symmetrical the pattern of effects was.

### RSA effects in regions of interest

These analyses investigated effects in our set of targeted ROIs that are known to be key nodes in the brain’s semantic network (Binder et al., 2009; Lambon Ralph et al., 2017). Correlations between semantic and neural DSMs for each ROI are shown in Figure 3C. These are consistent with the widespread and bilateral effects seen in the searchlight analyses. For the listening task, semantic similarity predicted neural similarity in all regions except IFG and right AG (at FDR-corrected *p* < 0.05). For the speaking task, all regions showed significant effects with the exception of vATL and right AG. In the cross-task analysis, significant correlations were observed in all regions except vATL. As with the searchlight results, in almost all cases semantic DSMs predicted neural similarity to a similar degree in the left and right hemispheres. Direct comparisons of the effects in left and right-hemisphere homologues revealed only two significant differences. In the production task, the correlation was significantly stronger in left vATL compared with right vATL. In the cross-task analysis, a significant difference was found in pMTG, but with *right* pMTG showing the stronger effect. Thus, while our univariate ROI analysis showed clearly that semantic regions in the left-hemisphere showed a stronger BOLD response during discourse processing, here we found that activation patterns in both hemispheres tracked the semantic content of discourse to a similar degree.

Results of control ROI analyses, using the quantity DSMs as the predictor, are provided in Supplementary Figure 5. A much more limited set of correlations was observed here. In the listening task, quantity DSMs predicted neural patterns in some regions, predominantly in the temporal lobes (left and right lATL, left vATL and right AG). In the speaking task, only one region (left AG) showed a correlation above zero, and only right IFG showed a cross-task effect. These results suggest that neural patterns in semantic brain regions were primarily sensitive to the content of discourse and not to the number of words processed in each segment.

### Comparison of similarity structure in different brain regions

The final analysis explored the degree to which different parts of the semantic network represent the content of discourse similarly. Rather than comparing each region’s neural DSM to a semantic DSM, here we directly compared the neural DSMs of different ROIs with each other (see Figure 1D). The correlations between ROIs for each analysis are shown in the top panel of Figure 6. All of the correlations were positive. This indicates consistency in neural responses across the brain: pairs of speech segments that elicited similar activation patterns in one ROI tended to elicit similar patterns in all of the other ROIs. In general, the strongest correlations between ROIs were observed in the listening task, with less convergence in the speaking task and the cross-task analysis. Strong correlations also tended to be observed between left and right homologue regions (visible as a diagonal line in the lower-right portion of each correlation plot). Correlations between left and right AG were particularly strong. The strong cross-hemispheric coupling in activation is also evident in the hierarchical cluster plots that group ROIs by similarity in their DSMs. In every case, left and right homologues were most similar to one another. This suggests a degree of functional association between left and right-hemisphere regions, despite the fact that the left hemisphere regions consistently showed stronger BOLD responses in the univariate analysis. Otherwise, the three cluster analyses showed broadly similar relationships between regions, with the pMTG and AG most strongly related and also clustered with the IFG, while ATL regions showed more distinct neural patterns.

**Figure 6:**
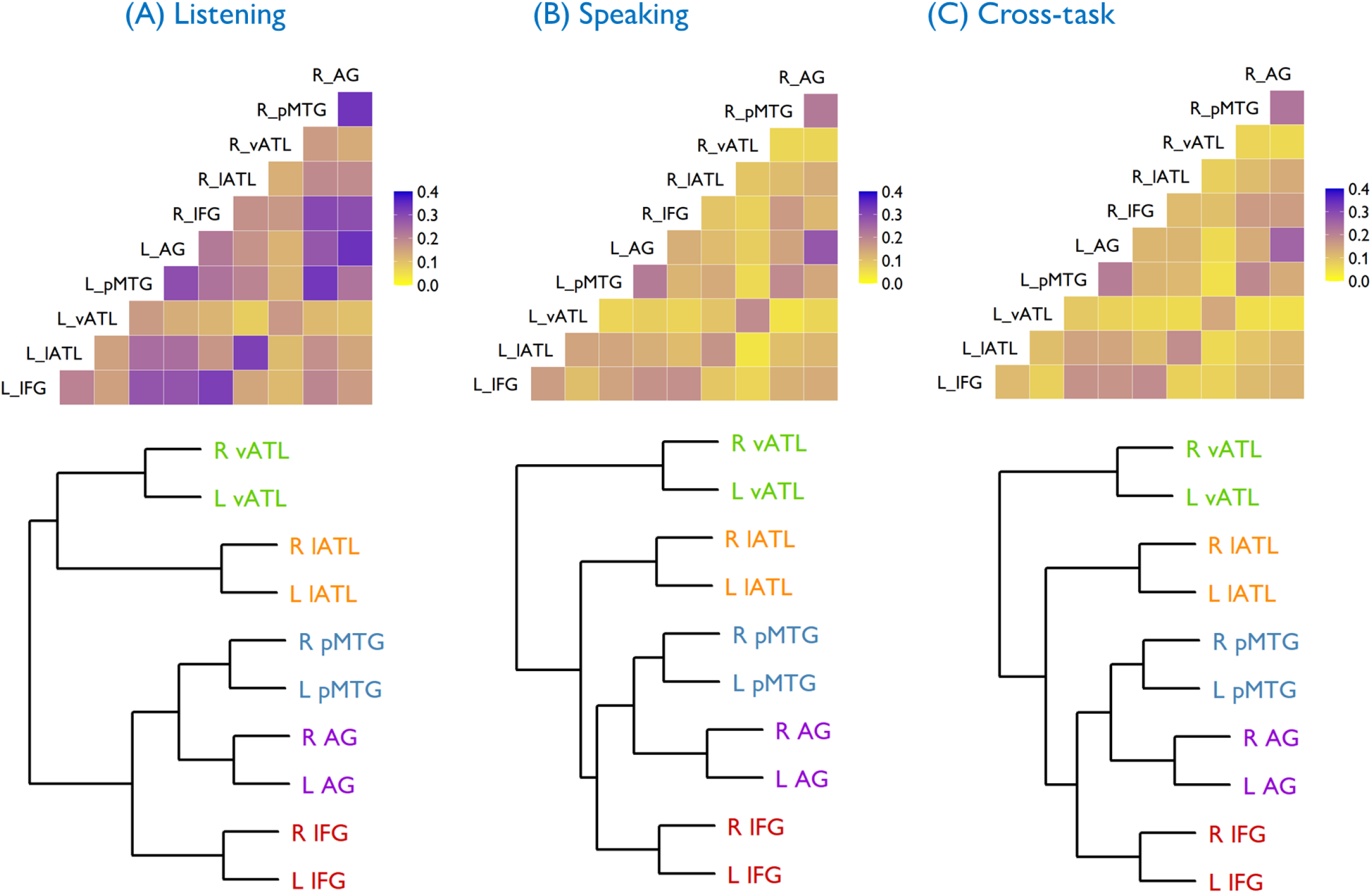
Relationships between neural DSMs. This analysis tested the correlations between neural DSMs obtained in different regions. Top panel shows correlations between neural DSMs for each analysis. Strong correlations indicate that neural DSMs were more similar to one another. Bottom panel shows hierarchical cluster analysis performed using the correlation data. The plots cluster regions according to how similar their neural DSMs are

## Discussion

Recent years have seen an explosion in investigations of how language content is represented in the brain, driven by advances in decoding and pattern analysis methods allied with more naturalistic fMRI paradigms (Dehghani et al., 2017; Huth et al., 2016; Schrimpf et al., 2021; Wehbe et al., 2014; Zhang et al., 2020). Crucially, however, prior studies have focused almost exclusively on how language is encoded during comprehension tasks, rather than when people produce their own discourse. Here, for the first time, we used a model of semantic discourse content to predict fMRI activation patterns during the *production* of language, comparing this to comprehension-based patterns in the same individuals. We imaged participants while they produced and listened to passages of discourse in response to topic prompts and we used RSA to relate neural similarities between different passages to similarity in their semantic content, using a vector-based model of distributional semantics (Landauer & Dumais, 1997). When participants produced discourse with similar content, they showed similar patterns of neural activity in a wide range of frontal, temporal and parietal sites, which were more extensive than those observed during comprehension. Moreover, semantic similarity also predicted neural similarities when comparing between the two language tasks, showing that concepts activated during production and comprehension share similar neural representations. Most importantly, the regions that showed semantic coding of discourse content in both production and comprehension were strikingly bilateral, suggesting a greater role for the right hemisphere in discourse processing than simple activation analyses have indicated.

One major contribution of the present work is to demonstrate similar neural coding of semantic content during production and comprehension of high-level discourse. Previous studies have shown inter-subject correlations when comparing the activation of a speaker and a comprehender (Liu et al., 2017; Silbert et al., 2014; Stephens et al., 2010), and have demonstrated that this coupling is greatest when both participants understand narratives in the same way (Heidlmayr, Weber, Takashima, & Hagoort, 2020). The present work makes two important new contributions. First, we show that similar neural patterns for production and comprehension occur within the same individuals as they switch between language tasks. Second, we show that comprehension-production similarities can be predicted by a distributional model of semantics, with semantically similar spoken/heard discourse passages eliciting more similar activation patterns. Alignment between comprehension and production is predicted by a range of theories which propose that higher levels of language processing are largely shared between production and comprehension (Hagoort, 2013; Kintsch & Vandijk, 1978; Levelt et al., 1999). Potential benefits of this shared cognitive machinery include the ability of the production system to make forward predictions that aid comprehension (Dell & Chang, 2014; Pickering & Garrod, 2013) and the ability for interlocutors to align their mental frameworks during conversation (Pickering & Garrod, 2004).

Semantic effects that crossed between language tasks were observed in a wide range of brain regions. Prominent among these were key nodes of the default-mode network: inferior parietal cortices (AG), posterior cingulate and medial prefrontal regions. The role of DMN regions in semantic processing is debated. While some authors have emphasised the similarities between the DMN and areas engaged by semantic tasks (Binder & Desai, 2011; Binder et al., 1999), others have argued for a greater distinction between these two systems (Humphreys, Hoffman, Visser, Binney, & Lambon Ralph, 2015). One complicating factor is that DMN regions frequently deactivate as tasks increase in difficulty (Mckiernan, Kaufman, Kucera-Thompson, & Binder, 2003). Activation differences between tasks can therefore be caused by uncontrolled differences in task demands. Because of this, deactivation does not necessarily imply that DMN regions are not participating in a cognitive process; for example, the posterior cingulate increases its connectivity with semantic regions during semantic tasks, while simultaneously showing deactivation effects (Krieger-Redwood et al., 2016). The RSA approach we employed here circumvents these issues by testing whether activation correlates reliably with semantic structure, rather than contrasting activity between conditions. These analyses suggest that a range of DMN sites are involved in the processing of linguistic meaning, at least at the level of discourse.

So, what is the functional role of these regions? Neuroimaging evidence implicates DMN regions in the construction of mental models that represent the details of events (situation models) in a range of contexts, including movie-watching (Baldassano et al., 2018), story-listening and reading (Ferstl, Neumann, Bogler, & von Cramon, 2008; Heidlmayr et al., 2020; Silbert et al., 2014), memory retrieval (Ranganath & Ritchey, 2012) and in social interactions (Yeshurun et al., 2021). Consistent with these findings, our previous work showed increased activation in this network when participants produced or heard discourse that was low in coherence, and thus was harder to construct a situation model for (Morales et al., 2022). The present results provide more direct evidence for the role of DMN regions in representing discourse content, by showing that they represent the semantics of discourse consistently across language processing modes.

With respect to AG in particular, our position is consistent with other work implicating this region in combinatorial semantic processing (i.e., computing the meaning of multi-word phrases) and event semantics (Binder & Desai, 2011; Mirman, Landrigan, & Britt, 2017; Price, Bonner, Peelle, & Grossman, 2015). In addition to semantic tasks, AG is activated by a number of other cognitive domains including autobiographical and episodic memory retrieval (Rugg & King, 2018), social cognition (Van Overwalle, 2009) and numerical processing (Sokolowski, Matejko, & Ansari, 2023). This has led to integrative accounts proposing that AG is involved in integrating and buffering multi-modal information over time (Humphreys, Lambon Ralph, & Simons, 2021; Seghier, 2013). These integrative accounts imply that AG is critical in constructing the underlying mental models that people use to both generate and understand discourse. Although there is evidence of specialisation for different cognitive domains across the inferior parietal region (Humphreys & Lambon Ralph, 2015; Seghier, Fagan, & Price, 2010), we found robust correlations of discourse content across the region. This is consistent with the idea that when people construct narratives they draw on a rich blend of verbal semantic knowledge, general world knowledge, specific personal experiences and understanding of social interactions.

We also found cross-task semantic effects in ATL regions. Though parts of the ATL are frequently identified as belonging to the default-mode network, theories of semantic cognition typically ascribe a different role to this region compared to other default-mode network regions like AG (Farahibozorg, Henson, Woollams, & Hauk, 2019; Humphreys et al., 2015; Humphreys et al., 2021; Mirman et al., 2017). While AG is thought to be involved in combining concepts to represent events and situations (Binder & Desai, 2011; Mirman et al., 2017; Price et al., 2015), theories of ATL function focus on its role in coding conceptual structure at the single-word and concept level (Lambon Ralph et al., 2017; Patterson et al., 2007). Our data support this idea that ATL is functionally distinct from AG, since cluster analyses indicated a strong separation between ATL similarity patterns and those of the other semantic regions we studied (including AG but also pMTG and IFG, which we will come to shortly). Nevertheless, there was some evidence that the ATLs were sensitive to the semantic content of speech. Lateral ATLs encoded semantic similarity during both conditions and in the cross-task analysis. The ventral part of the ATL was sensitive to semantic similarity only during speech comprehension. The lack of vATL effects during speech production may be due to this region’s high susceptibility to fMRI signal dropout (Ojemann et al., 1997).

ROI analyses also revealed effects of semantic similarity in the posterior temporal lobes (pMTG) and prefrontal cortices (IFG). pMTG has been implicated in the semantics of events and actions (Davey et al., 2016; Liljeström et al., 2008; Watson, Cardillo, Ianni, & Chatterjee, 2013), processes which are likely to be critical when understanding or generating narrative discourse. In addition, both pMTG and IFG form part of the semantic control network: a set of regions which show increased engagement when semantic processing requires high levels of cognitive control (Jackson, 2021; Noonan, Jefferies, Visser, & Lambon Ralph, 2013). Functions of this network include supporting retrieval of context-relevant semantic information as well as selection processes that arbitrate between multiple competing concepts to ensure contextually relevant information is attended to (Lambon Ralph et al., 2017). Semantic RSA effects in these regions might therefore reflect systematic variations in the cognitive control demands of different discourse topics.

At the whole-brain level, the most striking finding was the widespread right-hemisphere coding of discourse content. This occurred even though right-hemisphere regions were not activated to the same extent as left-hemisphere regions and even, in the case of production, were often deactivated relative to baseline conditions or to rest. The cross-ROI analysis further demonstrated that semantic content evoked similar activation patterns across left and right homologues of semantic processing regions. Thus, left and right-hemisphere regions appear to track the semantics of discourse in a similar way. This is an important general observation for neuroimaging studies because it suggests that regions may encode task-relevant information, even without showing increases in BOLD signal in a traditional subtraction design.

At the same time, the disparities in activation levels suggest that the left hemisphere was more actively engaged than the right when processing meaningful discourse. One interpretation of this is that discourse content is represented bilaterally but that the left hemisphere dominates processing under normal conditions. This redundancy in representation may provide the system with some resilience in the event of damage to the left hemisphere (Rice, Hoffman, & Lambon Ralph, 2015). Indeed, semantic-related activation in the right hemisphere increases when dominant left-hemisphere regions are impaired: this pattern has been observed following brain stimulation (Jung & Lambon Ralph, 2016) andsurgical resection of the left ATL (Rice et al., 2018), as well as in the context of healthy ageing (Hoffman & Morcom, 2018).

The bilateral RSA results are also consistent with the view that the right hemisphere makes specific contributions to understanding natural language, such as understanding metaphors and jokes (Marinkovic et al., 2011; Rapp, Mutschler, & Erb, 2012), making inferences (Mason & Just, 2004) and comprehending narrative structures (Knutson, Wood, & Grafman, 2004). One particular framework suggests that semantic processing occurs bilaterally, with each hemisphere undertaking its own type of neurocomputation but working interactively with the other (Jung-Beeman, 2005). The left hemisphere shows fine coding, rapidly activating a network of strongly linked semantic associates; while the right hemisphere encodes more coarse associations, and as such is sensitive to more distant semantic relations and broader conceptual meanings. This coarse coding is thought to be more important for capturing the nuanced meanings contained in more complex language acts, such as extended narratives. Indeed, our findings are consistent with previous studies that have implicated a bilateral network in comprehension of extended, naturalistic speech (e.g., Huth et al., 2016). Importantly, however, we have shown that this right-hemisphere contribution is also present in discourse production. While few neuroimaging studies have investigated language production, lesion studies indicate that patients with right-hemisphere damage produce disorganised and poorly structured narratives, suggesting a right-hemisphere role in discourse planning processes (Bartels-Tobin & Hinckley, 2005; Davis, O’Neil-Pirozzi, & Coon, 1997; Marini, Carlomagno, Caltagirone, & Nocentini, 2005). Studies of ATL function also point to a bilateral pattern of engagement in lexical-semantic processing (Rice et al., 2015). Our results are broadly in line with these findings in suggesting that right-hemisphere homologues of semantic regions code the semantic properties of discourse during production as well as comprehension.

We end by considering ways in which the present approach could be extended in future work. First, our semantic model uses information about patterns of word usage in natural language to determine similarity in meaning. Though such distributional approaches can very accurately mimic human judgements of semantic relatedness (Pereira, Gershman, Ritter, & Botvinick, 2016), they have been criticized on the grounds that they do not capture perceptual properties of objects such as their size, shape or colour (Bruffaerts et al., 2019; Glenberg & Mehta, 2008; Glenberg & Robertson, 2000). Future studies may benefit from the use of more complex semantic models that make use of both lexical co-occurrence and experiential attributes (Andrews, Vigliocco, & Vinson, 2009; Davis & Yee, 2021; Fernandino, Tong, Conant, Humphries, & Binder, 2022; Hoffman, McClelland, & Lambon Ralph, 2018b). Applying such an approach to discourse (cf. single-word comprehension) will present significant challenges, as discourse contains descriptions of complex situations whose perceptual characteristics cannot easily be predicted from their lexical constituents. Second, conceptual representations are dynamic and are shaped by the individual’s personal context, in terms of their past experiences, individual processing preferences and abilities (Yee & Thompson-Schill, 2016). We controlled for these individual differences by directly comparing comprehension and production in the same individuals (as opposed to previous studies which compared speaking and listening in different people). Nevertheless, the degree to which people’s discourse representations vary in their content and neural instantiation remains an important question for future work.

It is also important to bear in mind that the act of producing speech inherently entails auditory processing of one’s own utterances. There is evidence that auditory cortical responses to self-produced speech are suppressed compared with other-produced speech (Heinks-Maldonado, Mathalon, Gray, & Ford, 2005). Indeed we observed less superior temporal activation in our speaking condition compared to the listening condition. Nevertheless, participants did hear their own utterances and this auditory processing may have contributed to the activation of semantic knowledge in the speaking task. This in turn could have contributed to the common coding we observed in the cross-task analysis.

Finally, our distributional semantic model uses a “bag of words” approach: semantic representations are computed by averaging over all the words in passage, without taking word order into account. Semantic similarity scores calculated in this way are strongly correlated with human ratings of passage similarity (McNamara, Cai, & Louwerse, 2007; Stone, Dennis, & Kwantes, 2011). Nevertheless, this approach cannot capture information about roles and relations between discourse elements (e.g., the distinction between “a dog chases a person” and “a person chases a dog”). More advanced transformer-based language models (Devlin, Chang, Lee, & Toutanova, 2018) may be better able to capture these aspects of discourse structure.

To conclude, we have used a distributional model of semantics to predict neural similarity patterns during discourse processing. We have shown that a broad set of brain regions code language content during speech production in a similar way to when hearing someone else’s speech, suggesting common coding across different language processing modes. We believe that this work can stimulate further insights into how the semantic systems of the brain drive the generation, as well as understanding, of language.

### Data and Code Availability

Data and code associated with this study are available at: https://osf.io/s3z8y/. Unthresholded activation and correlation maps are archived at: https://neurovault.org/collections/12792/.

## Supplementary Materials

### List of prompts used in production and comprehension tasks

Speaking

1. Describe how you would make a cup of tea or coffee.
2. Why are some people concerned about climate change?
3. Why is it important to have a balanced diet?
4. Describe the steps you’d take to order-in food.
5. Describe a typical visit to a grocery store.
6. How would you prepare to go on holiday?
7. Do you think the internet has improved people’s lives?
8. What would you recommend doing during a job interview?
9. What do people usually do on New Year’s Eve in the UK?
10. What sorts of things do people do to cope with stress?
11. What sort of things does a teacher do when at work?
12. What sort of things usually happen at a wedding?

Listening

1. What would it be like to live in Antarctica?
2. What happens when a storm is forecast in the UK?
3. What do the police do when a crime has been committed?
4. Which is your favourite season and why?
5. What do you like or dislike about Christmas?
6. Do you think it’s a good idea to send people to live on Mars?
7. Why do people come to Scotland on holiday?
8. What sort of things do you have to do to look after a dog?
9. Describe a typical visit to a restaurant.
10. What are the advantages and disadvantages of going to university?
11. What do people usually do when getting ready for work in the morning?
12. Describe the steps you would need to take if going somewhere by train.

### Example sample presented in the listening task

#### Prom pt: What would it be like to live in Antarctica?

“I don’t know whether Antarctica is north or south but I imagine that it’s cold. It’s one or the other. I don’t know if there are penguins there. But there would be penguins around, I guess. If there were penguins there they would in the Antarctic with you. You’d probably be living in an igloo, riding reindeers and stuff and killing seals to survive and wearing polar bear hide or something weird. I don’t know. You’d probably be pretty chilly all the time, so you’d have to layer up a lot. Beware of killer whales that burst through the ice and eat you in your sleep. You’ve got to really just be aware of the dangers of nature. Also snowstorms. They can pretty killer. If you get stuck in a snowstorm you’re pretty screwed because they can be pretty harsh”

### Example sample produced in the speaking task

#### Prom pt: What would you recom m end doing during a job interview?

“I would recommend during a job interview to come across as professional and know what you’re talking about. I think preparation for a job interview is key, so learn about the company, why you want to work there, their values and strengths that you could bring to the company and what the company can bring to you. I think it’s very important that you come across as approachable and true to your word and morals. I think it’s very important for people who are interviewing people to really trust their interviewees and to get a good sense of who they are and what they believe in. I think it’s really important to be honest about what you can do, but obviously, be confident in yourself.”

## Supplementary Figures

**Supplementary Figure 1:**
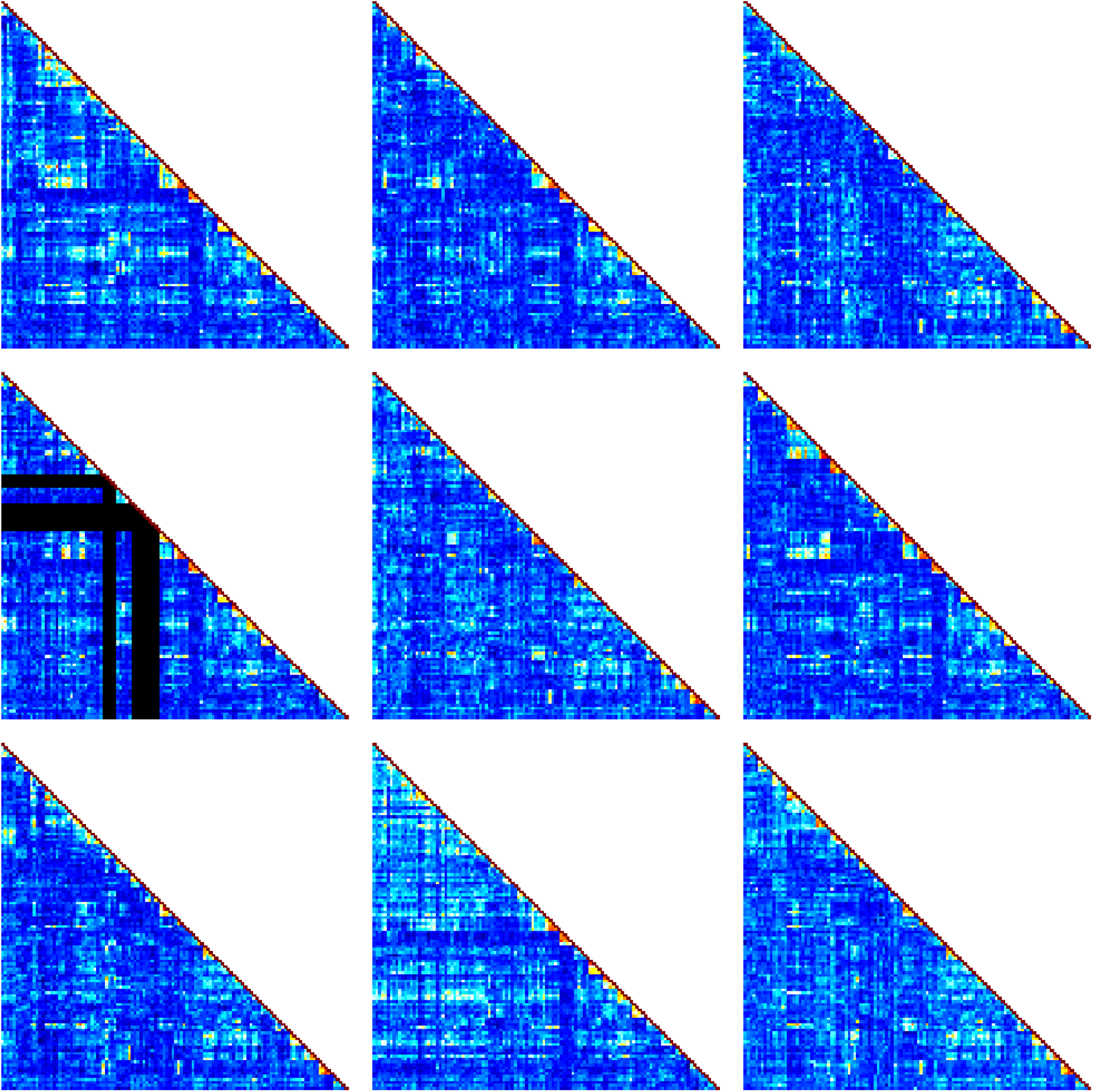
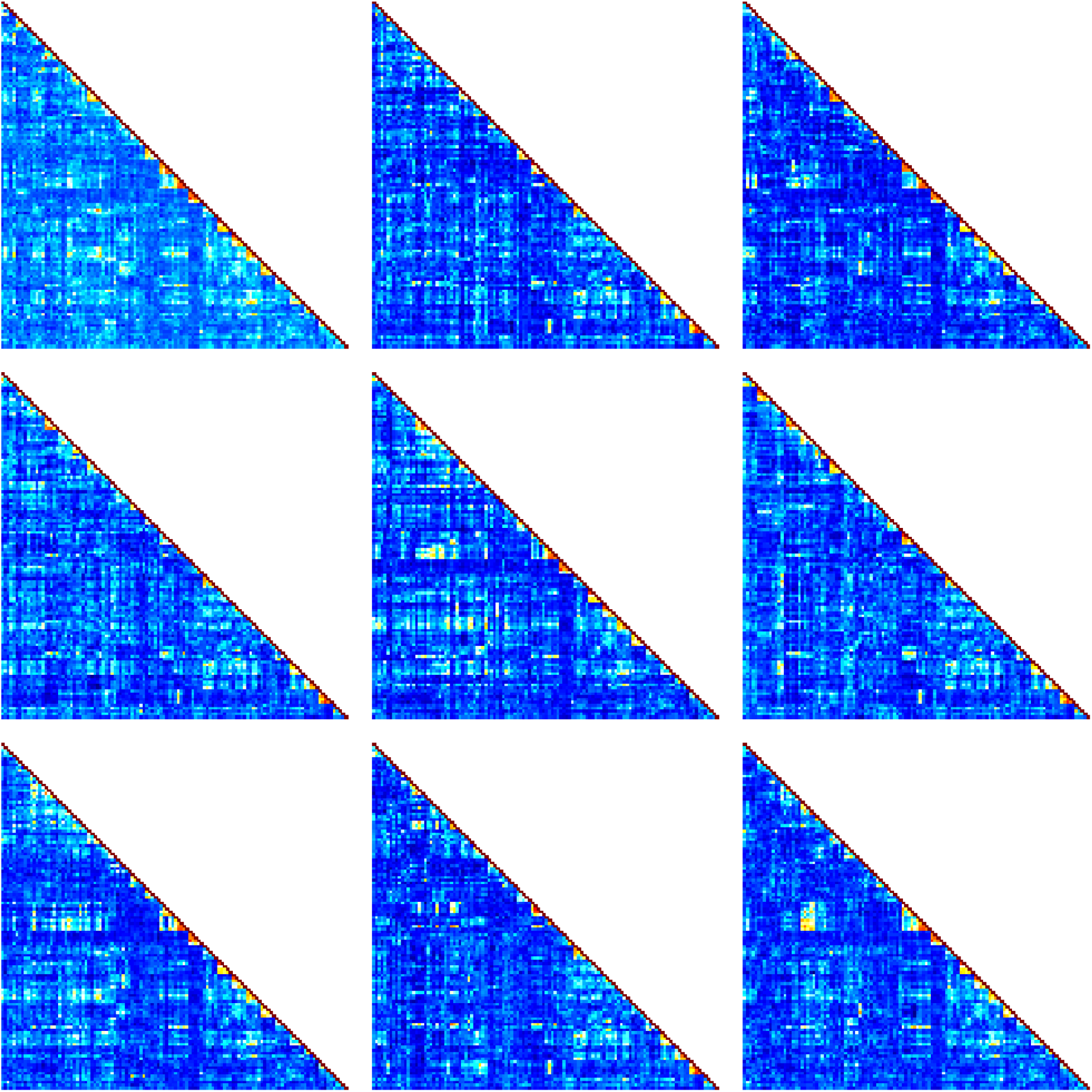
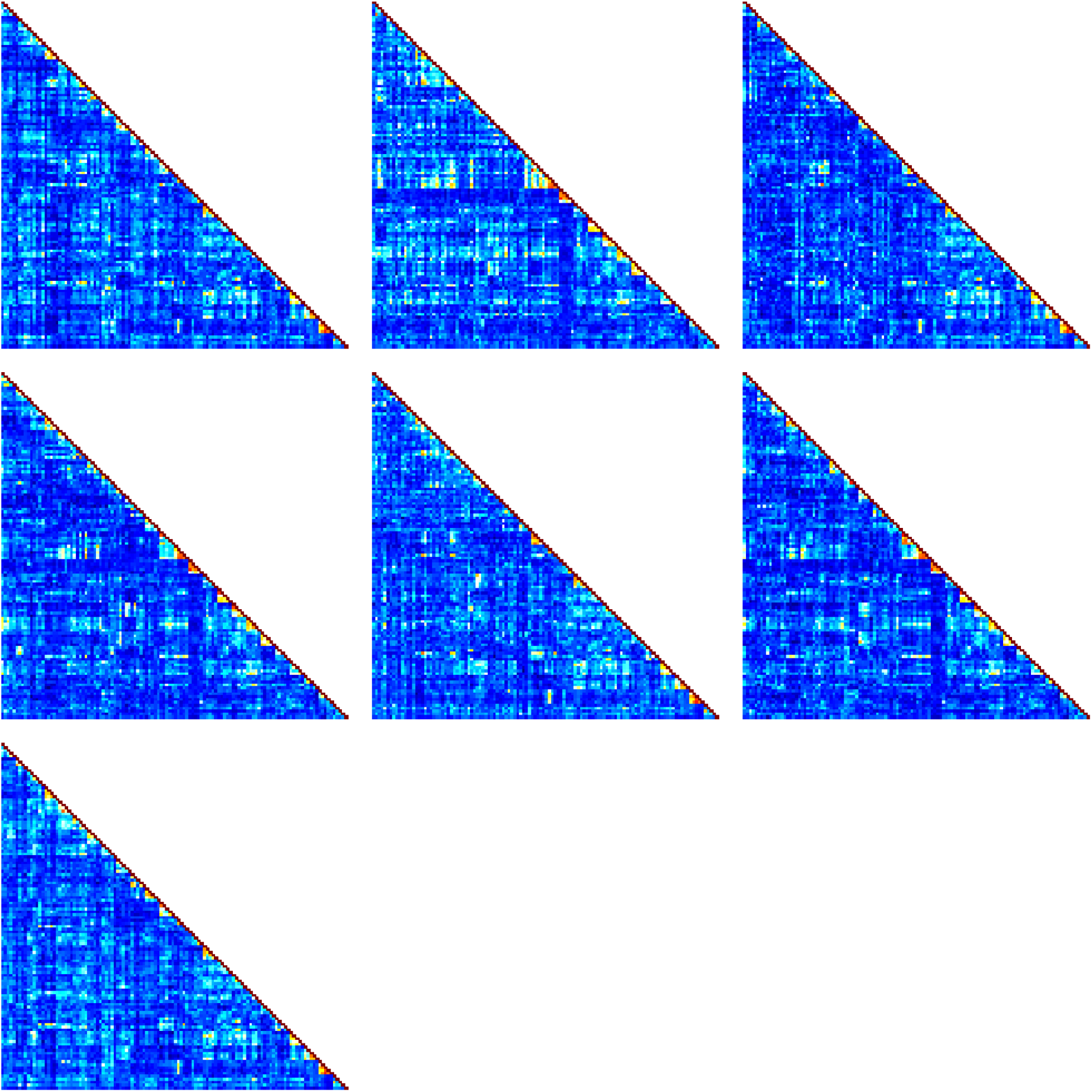
Semantic DSMs for all participants. We present the semantic dis-similarity matrix for each participant on the following three pages. There is some missing data (shown in black) for one participant as audio recording was unsuccessful for three of their speaking topics.

**Supplementary Figure 2:**
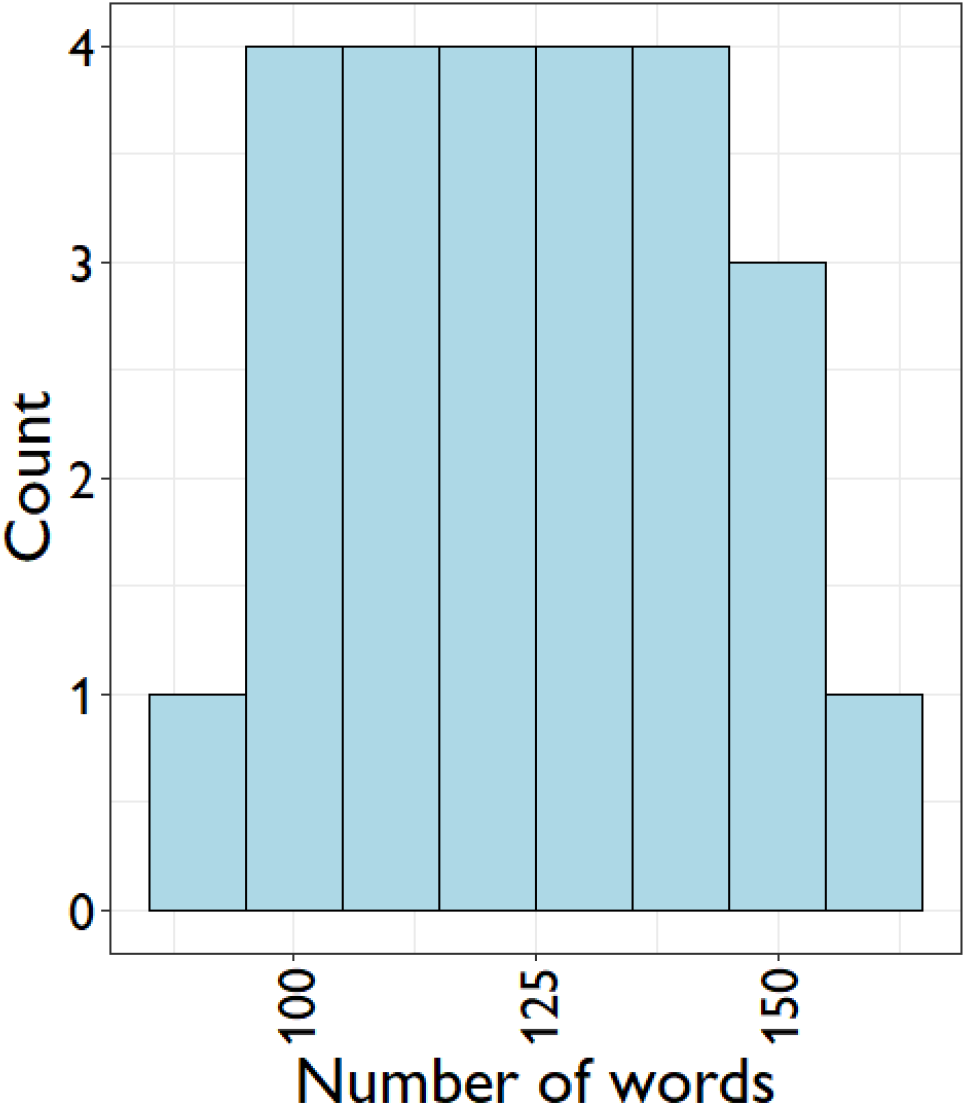
Number of words produced by participants in speaking task. Histogram shows the distribution over participants of the mean number of words produced per topic.

**Supplementary Figure 3:**
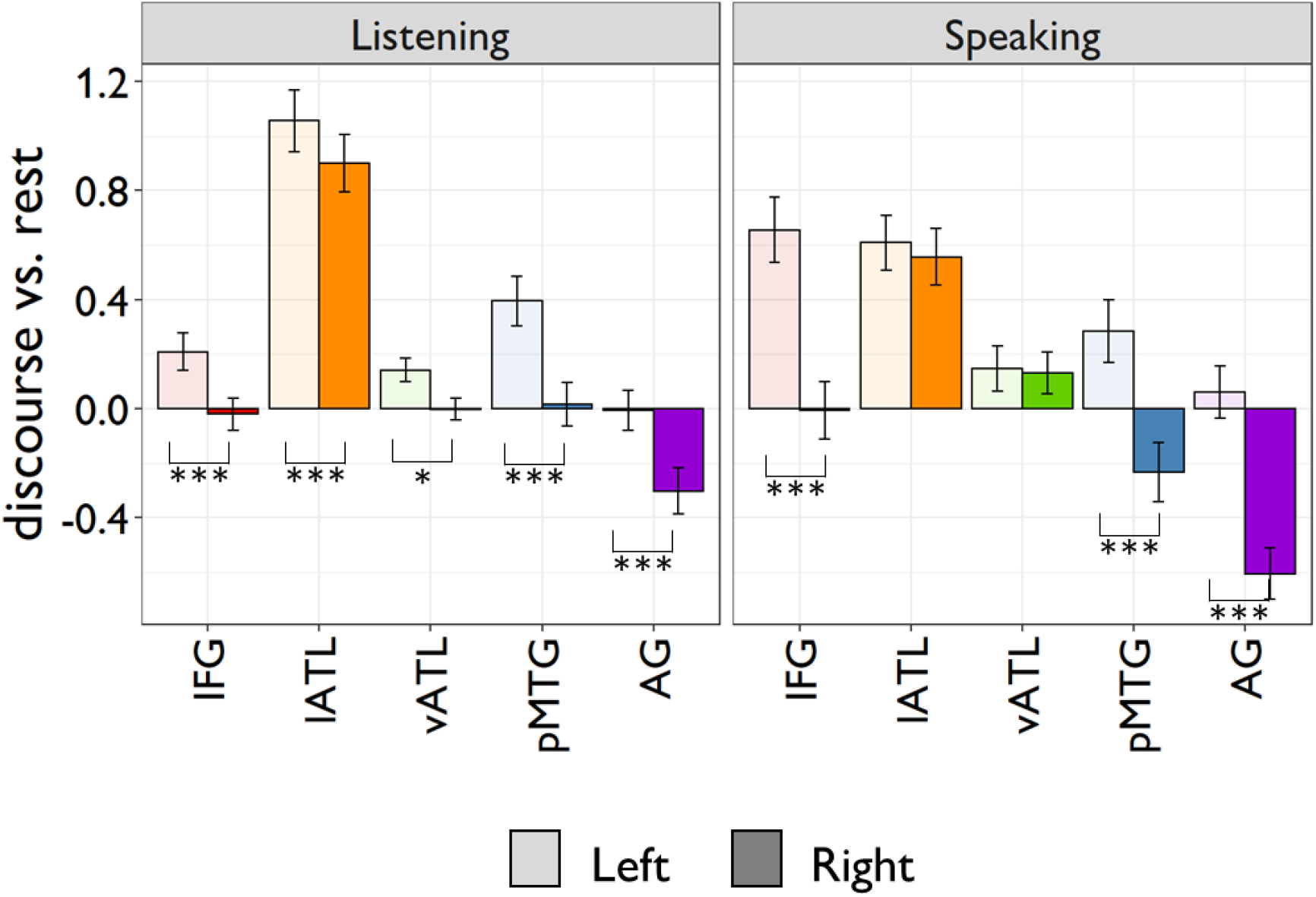
Effects of discourse relative to rest in semantic regions. Left hemisphere regions shown in pale tones and right hemisphere in solid tones. Asterisks below the x-axis indicate significant hemispheric differences. * = p < 0.05; ** = p < 0.01; *** = p < 0.001, all FDR-corrected. Error bars show 1 SEM. A 2 x 2 x 5 (task x hemisphere x ROI) ANOVA performed on these data revealed main effects of ROI (F(4,96) = 75.6, p < .001) and hemisphere (F(1,24) = 60.2, p < .001; left > right) but no effect of task (F(1,24) = 0.8, p = .39). However, all of the interactions between these factors were significant (F > 5.4, p < .028).

**Supplementary Figure 4:**
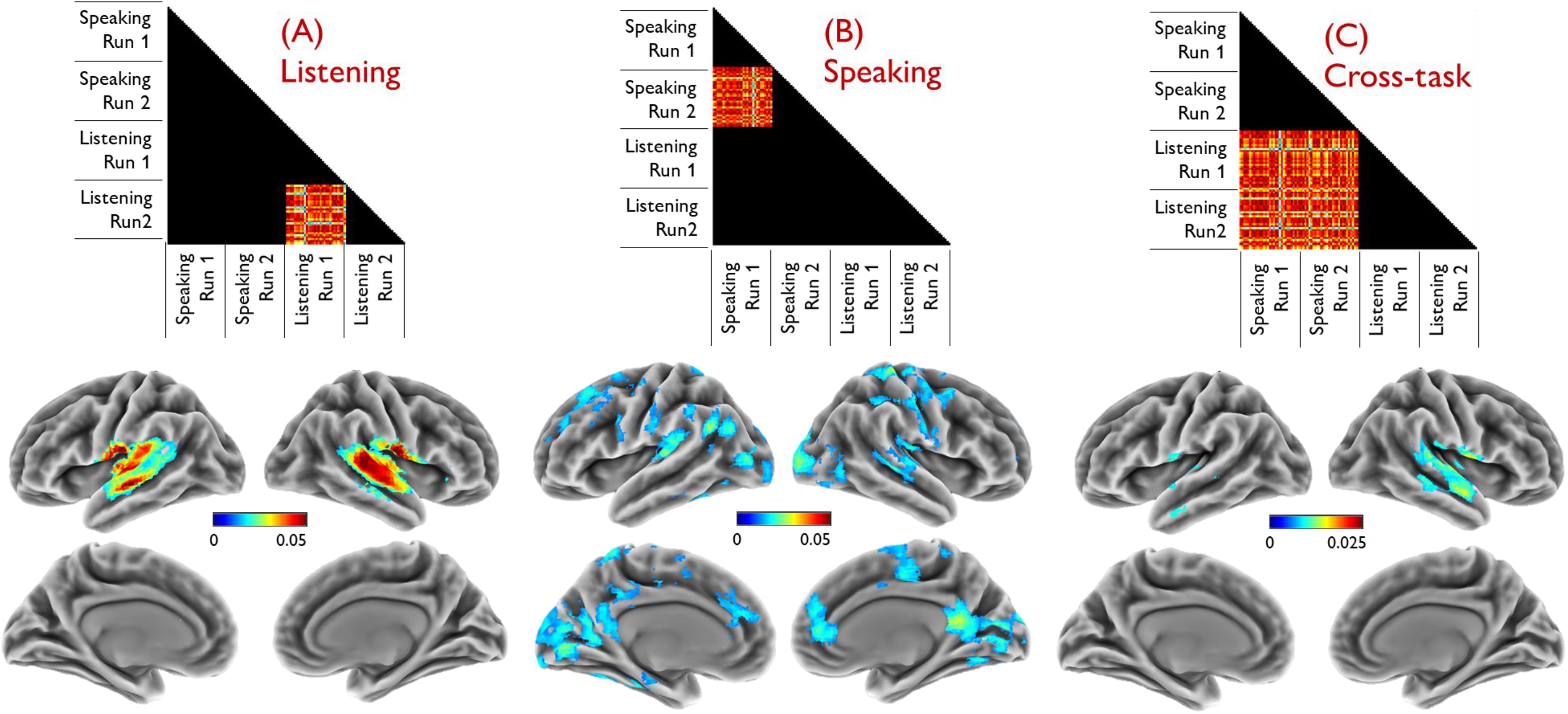
Searchlight analyses using the quantity DSM. Top panel indicates which speech parts of the DSMs were used in each analysis (between-run speech pairs in A and B; between-task pairs in C). Brain maps show regions where correlation between neural and quantity DSMs exceeded zero (at cluster-corrected p <0.05). Colour scales show the Fisher-transformed correlation coefficient.

**Supplementary Figure 5:**
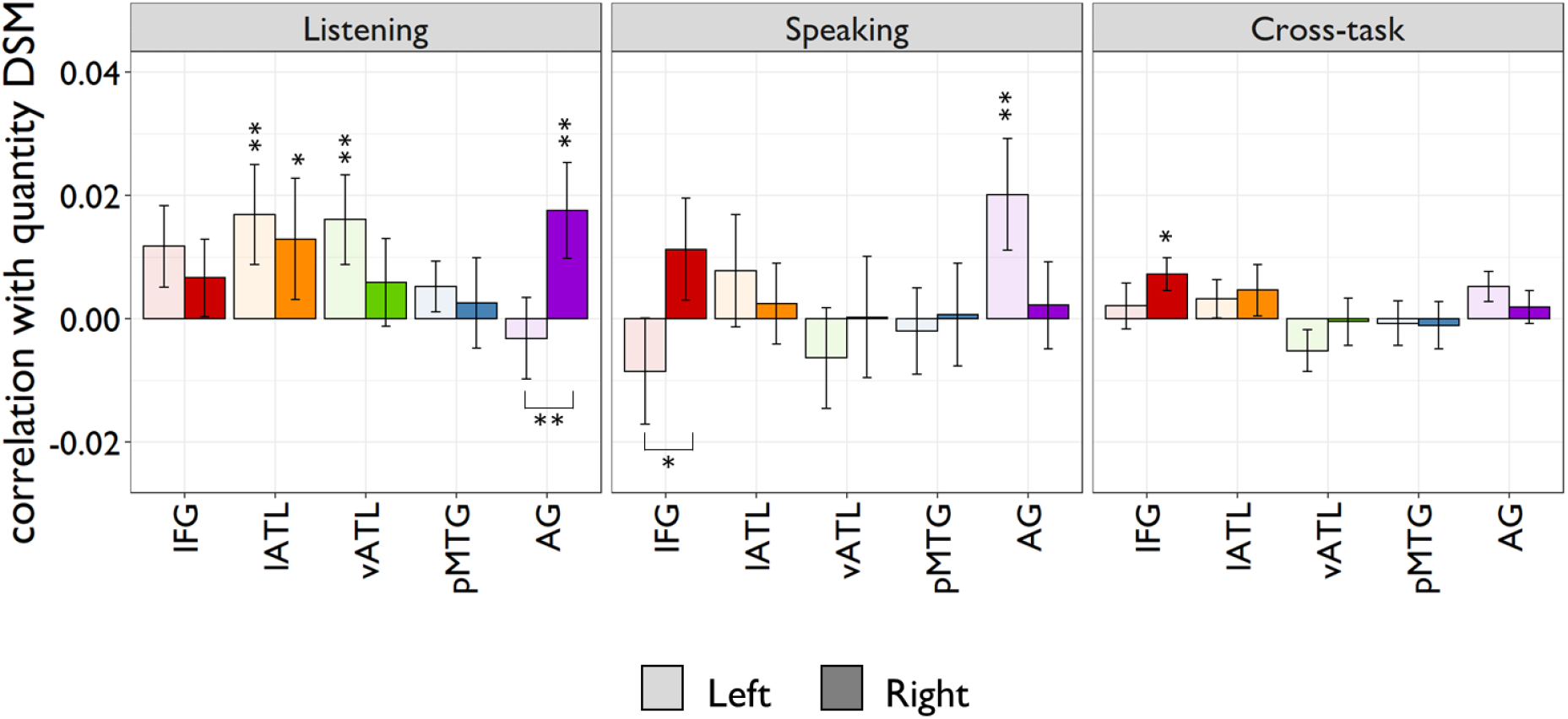
Region of interest results using the quantity DSMs. Left hemisphere regions shown in pale tones and right hemisphere in solid tones. Asterisks below the x-axis indicate significant hemispheric differences. Asterisks above the x-axis indicate correlations significantly greater than zero. * = p < 0.05; ** = p < 0.01; *** = p < 0.001, all FDR-corrected. Error bars show 1 SEM.

**Supplementary Table 1:** Peak activations for univariate analyses.

| *Contrast* | *Cluster extent* | *Location* | *t* | *x* | *y* | *z* |
| --- | --- | --- | --- | --- | --- | --- |
| Listening > Baseline | 1113 | R cerebellum | 10.91 | 21 | -81 | -39 |
|  |  | R lingual gyrus | 4.38 | 21 | -87 | -3 |
|  |  | R cerebellum | 3.27 | 30 | -69 | -57 |
|  | 8666 | L temporal pole | 9.51 | -48 | 15 | -30 |
|  |  | L IFG (pars triangularis) | 9.18 | -60 | 21 | 12 |
|  |  | L superior temporal gyrus | 8.88 | -51 | -6 | -15 |
|  |  | L middle frontal gyrus | 8.27 | -39 | 9 | 45 |
|  |  | L hippocampus | 8.24 | -30 | -12 | -15 |
|  |  | L superior frontal gyrus | 7.54 | -12 | 36 | 60 |
|  |  | L pMTG | 7.46 | -54 | -36 | -3 |
|  |  | L superior frontal gyrus | 6.98 | -12 | 57 | 39 |
|  |  | L IFG (pars orbitalis) | 6.54 | -42 | 36 | -9 |
|  |  | R temporal pole | 6.49 | 48 | 9 | -27 |
|  |  | L angular gyrus | 6.47 | -45 | -54 | 24 |
|  |  | R amygdala | 5.70 | 30 | -6 | -21 |
|  |  | L superior frontal gyrus | 5.49 | -9 | 12 | 66 |
|  |  | R superior frontal gyrus | 5.25 | 12 | 54 | 42 |
|  |  | R pMTG | 5.20 | 48 | -36 | -6 |
|  |  | L fusiform gyrus | 4.54 | -39 | -45 | -18 |
|  |  | L precentral gyrus | 4.44 | -12 | -21 | 57 |
|  |  | R superior parietal lobule | 4.37 | 21 | -45 | 63 |
|  |  | Brainstem | 4.32 | -6 | -30 | -6 |
|  |  | R superior frontal gyrus | 4.29 | 9 | -12 | 63 |
| Speaking > Baseline | 12534 | R cerebellum | 12.92 | 18 | -81 | -42 |
|  |  | L IFG (pars triangularis) | 10.09 | -45 | 27 | 3 |
|  |  | L superior frontal gyrus | 9.89 | -9 | 15 | 60 |
|  |  | L angular gyrus | 9.83 | -42 | -57 | 27 |
|  |  | L superior frontal gyrus | 9.68 | -12 | 63 | 24 |
|  |  | L superior frontal gyrus | 9.31 | -12 | 48 | 39 |
|  |  | L middle frontal gyrus | 9.22 | -39 | 9 | 60 |
|  |  | R cerebellum | 8.15 | 42 | -69 | -48 |
|  |  | L caudate nucleus | 8.02 | -15 | 3 | 21 |
|  |  | R caudate nucleus | 7.75 | 12 | 0 | 21 |
|  |  | R cerebellum | 7.75 | 3 | -54 | -42 |
|  |  | R cerebellum | 7.70 | 33 | -60 | -30 |
|  |  | L middle temporal gyrus | 7.13 | -57 | -12 | -15 |
|  |  | L calcarine gyrus | 7.13 | -6 | -57 | 9 |
|  |  | L frontal pole | 6.67 | -3 | 63 | -12 |
|  |  | R calcarine gyrus | 6.47 | 24 | -51 | 9 |
|  |  | Cerebellar vermis | 6.43 | 0 | -48 | -18 |
|  |  | L middle frontal gyrus | 5.85 | -51 | 21 | 27 |
|  |  | R calcarine gyrus | 5.62 | 24 | -72 | 12 |
|  |  | L frontal pole | 5.32 | -24 | 42 | 12 |
| Listening ><br>Speaking | 521 | R IFG (pars triangularis) | 5.50 | 51 | 12 | 27 |
|  |  | R middle frontal gyrus | 4.38 | 30 | 0 | 48 |
|  |  | R middle frontal gyrus | 3.83 | 51 | 15 | 48 |
|  |  | R IFG (pars triangularis) | 3.30 | 57 | 36 | 18 |
|  | 442 | L parahippocampal gyrus | 4.98 | -27 | 0 | -24 |
|  |  | L superior temporal gyrus | 4.68 | -51 | -6 | -12 |
|  |  | L temporal pole | 2.88 | -45 | 12 | -36 |
|  | 639 | R postcentral gyrus | 4.61 | 12 | -39 | 81 |
|  |  | L postcentral gyrus | 4.45 | -9 | -42 | 63 |
|  |  | L middle cingulate cortex | 4.26 | -12 | -24 | 51 |
|  |  | R superior parietal lobule | 3.77 | 15 | -54 | 66 |
|  |  | L postcentral gyrus | 3.57 | -30 | -36 | 48 |
|  |  | L postcentral gyrus | 3.53 | -24 | -36 | 75 |
|  |  | R middle cingulate cortex | 3.33 | 6 | -12 | 51 |
| Speaking ><br>Listening | 1360 | R cerebellum | 6.43 | 33 | -63 | -24 |
|  |  | R cerebellum | 5.20 | 33 | -63 | -48 |
|  |  | Cerebellar vermis | 4.37 | 0 | -63 | -30 |
|  |  | R cerebellum | 4.22 | 21 | -36 | -42 |
|  |  | R cerebellum | 3.58 | 3 | -42 | -21 |
|  |  | R cerebellum | 3.36 | 12 | -60 | -3 |
|  |  | R cerebellum | 3.31 | 9 | -84 | -39 |
|  | 920 | L superior frontal gyrus | 5.27 | -9 | 12 | 60 |
|  |  | L superior frontal gyrus | 4.81 | -21 | 33 | 57 |
|  |  | L superior frontal gyrus | 4.48 | -12 | 45 | 36 |
|  |  | L middle frontal gyrus | 4.43 | -36 | 9 | 54 |
|  |  | L anterior cingulate gyrus | 3.50 | -6 | 24 | 39 |
|  | 471 | L calcarine gyrus | 4.57 | -18 | -81 | 9 |
|  |  | L Lingual gyrus | 4.47 | -12 | -69 | -6 |
|  |  | L parahippocampal gyrus | 3.57 | -33 | -42 | -6 |

Table shows co-ordinates in MNI space. L = left; R = right; IFG = inferior frontal gyrus; pMTG = posterior middle temporal gyrus.

**Supplementary Table 2:** Peak effects for representational similarity analyses.

| *Contrast* | *Cluster extent* | *Location* | *r* | *x* | *y* | *z* |
| --- | --- | --- | --- | --- | --- | --- |
| Listening | 1703 | R temporal pole | 0.037 | 51 | 24 | -21 |
|  |  | R inferior temporal gyrus | 0.030 | 45 | -3 | -30 |
|  |  | R superior temporal gyrus | 0.030 | 54 | -12 | -9 |
|  |  | R pMTG | 0.029 | 60 | -39 | 9 |
|  |  | R ventral temporal pole | 0.023 | 39 | 6 | -51 |
|  |  | R inferior temporal gyrus | 0.020 | 54 | -33 | -15 |
|  |  | R pallidum | 0.019 | 24 | -12 | 3 |
|  |  | R Precentral gyrus | 0.018 | 66 | 0 | 6 |
|  |  | R rolandic operculum | 0.016 | 51 | -30 | 27 |
|  | 188 | L superior frontal gyrus | 0.036 | -21 | 21 | 36 |
|  |  | L superior frontal gyrus | 0.023 | -21 | 30 | 60 |
|  | 4012 | R calcarine gyrus | 0.034 | 18 | -78 | 21 |
|  |  | L cuneus | 0.031 | -12 | -84 | 33 |
|  |  | L posterior cingulate cortex | 0.028 | -6 | -45 | 36 |
|  |  | L supramarginal gyrus | 0.025 | -45 | -48 | 24 |
|  |  | R precuneus | 0.025 | 9 | -60 | 45 |
|  |  | R angular gyrus | 0.024 | 51 | -72 | 24 |
|  |  | L lingual gyrus | 0.024 | -21 | -66 | 0 |
|  |  | L precuneus | 0.023 | -6 | -75 | 60 |
|  |  | Occipital pole | 0.022 | 0 | -96 | 21 |
|  |  | R lingual gyrus | 0.022 | 27 | -57 | -3 |
|  | 1109 | L cerebellum | 0.032 | -27 | -72 | -51 |
|  |  | R cerebellum | 0.030 | 18 | -63 | -39 |
|  |  | L cerebellum | 0.027 | -6 | -51 | -42 |
|  |  | Brainstem | 0.024 | -3 | -36 | -60 |
|  |  | R cerebellum | 0.022 | 24 | -87 | -45 |
|  | 269 | L middle temporal gyrus | 0.027 | -54 | -9 | -21 |
|  |  | L temporal pole | 0.018 | -57 | 12 | -27 |
|  | 284 | L basal ganglia | 0.026 | -15 | 9 | -9 |
|  |  | L middle frontal gyrus | 0.024 | -24 | 42 | 9 |
|  |  | L middle frontal gyrus | 0.021 | -39 | 57 | 18 |
|  | 103 | R middle frontal gyrus | 0.025 | 51 | 9 | 51 |
|  |  | R postcentral gyrus | 0.021 | 63 | -3 | 36 |
|  | 372 | R superior frontal gyrus | 0.024 | 21 | 27 | 36 |
|  |  | R superior frontal gyrus | 0.024 | 15 | 36 | 60 |
|  |  | R superior frontal gyrus | 0.022 | 21 | 54 | 36 |
|  | 7 | R orbitofrontal cortex | 0.023 | 9 | 51 | -27 |
|  | 33 | L anterior fusiform gyrus | 0.022 | -39 | -9 | -48 |
|  | 59 | L orbitofrontal cortex | 0.021 | -21 | 30 | -27 |
|  | 36 | Midbrain | 0.021 | 0 | -27 | -12 |
| Speaking | 15398 | L lateral occipital cortex | 0.050 | -30 | -78 | 30 |
|  |  | L pMTG | 0.043 | -51 | -51 | -3 |
|  |  | R lateral occipital cortex | 0.039 | 39 | -72 | 21 |
|  |  | R superior occipital gyrus | 0.036 | 27 | -78 | 42 |
|  |  | L IFG (pars triangularis) | 0.036 | -45 | 27 | 9 |
|  |  | R superior Parietal Lobule | 0.036 | 33 | -54 | 57 |
|  |  | R inferior temporal gyrus | 0.036 | 54 | -57 | -12 |
|  |  | L orbitofrontal gyrus | 0.035 | -24 | 33 | -6 |
|  |  | R middle frontal gyrus | 0.035 | 33 | -3 | 57 |
|  |  | L superior parietal lobule | 0.034 | -18 | -57 | 54 |
|  | 113 | L subgenual cingulate cortex | 0.021 | -3 | 24 | -12 |
|  | 21 | Midbrain | 0.018 | -3 | -18 | -6 |
|  | 11 | R supramarginal gyrus | 0.016 | 36 | -27 | 36 |
| Cross-Task | 26496 | L middle temporal gyrus | 0.023 | -54 | -9 | -21 |
|  |  | L Lateral occipital cortex | 0.023 | -27 | -87 | 42 |
|  |  | R temporal pole | 0.020 | 45 | 12 | -24 |
|  |  | R pMTG | 0.020 | 69 | -51 | -3 |
|  |  | R IFG (pars triangularis) | 0.019 | 48 | 33 | 12 |
|  |  | R middle frontal gyrus | 0.019 | 39 | 21 | 48 |
|  |  | L lingual gyrus | 0.019 | -27 | -66 | 3 |
|  |  | L inferior temporal gyrus | 0.018 | -45 | -51 | -6 |
|  |  | L IFG (pars triangularis) | 0.018 | -51 | 30 | 9 |
|  |  | R angular gyrus | 0.017 | 54 | -63 | 33 |
|  |  | R frontal pole | 0.017 | 21 | 63 | 12 |
|  |  | R subgenual cingulate cortex | 0.016 | 9 | 18 | -30 |
|  |  | L angular gyrus | 0.016 | -36 | -63 | 33 |
|  |  | L parahippocampal Gyrus | 0.015 | -27 | -27 | -21 |
|  |  | R fusiform gyrus | 0.015 | 39 | -36 | -15 |
|  |  | L middle frontal gyrus | 0.015 | -39 | 27 | 48 |
|  |  | R cerebellum | 0.015 | 21 | -60 | -18 |
|  |  | L lateral occipital cortex | 0.015 | -24 | -84 | 12 |
|  |  | L Superior frontal Gyrus | 0.015 | -15 | 33 | 33 |
|  |  | R inferior temporal gyrus | 0.014 | 54 | -18 | -21 |
|  |  | L orbitofrontal cortex | 0.014 | -24 | 36 | -9 |
|  |  | L frontal pole | 0.014 | -6 | 63 | 27 |
|  |  | R posterior cingulate cortex | 0.014 | 3 | -54 | 30 |
|  |  | R lateral occipital cortex | 0.014 | 36 | -69 | 21 |
|  |  | L orbitofrontal cortex | 0.014 | -18 | 30 | -30 |

Table shows co-ordinates in MNI space. L = left; R = right; IFG = inferior frontal gyrus; pMTG = posterior middle temporal gyrus.

